# Elucidation and functional characterization of natural sweetener biosynthesis in hortensia species

**DOI:** 10.1101/2024.10.09.617031

**Authors:** Goutham Padmakumar Sarala, Frauke Engel, Anja Hartmann, Nicolaus von Wirén, Mohammad-Reza Hajirezaei

## Abstract

Among the various coumarins found in Hydrangea species, the bioactive dihydroisocoumarin known as phyllodulcin (PD) stands out as a non-caloric, high-intensity sweetener, which is up to 800 times sweeter than sucrose. Additionally, PD possesses notable medicinal properties, including antifungal, anti-ulcer, and anti-inflammatory effects. PD also plays several plant-specific roles, such as defending against phytopathogens, responding to abiotic and biotic stresses and regulating hormones. However, the biological function and biosynthetic pathway of PD in Hydrangea remain unexplored. This study employs targeted metabolomic and transcriptomic approaches to uncover the regulatory mechanisms and identify potential candidate genes involved in the biosynthesis of PD. 14 out of 182 different Hydrangea accessions, exhibiting varying levels of PD, were selected for detailed analysis. Accessions of *H. macrophylla* with high PD levels displayed distinct metabolite profiles compared to those with low PD concentrations. Specifically, pathways involving caffeic acid, ferulic acid, and their derivatives, such as scopolin, scopoletin, esculetin, and fraxetin, were predominant in accessions with low PD concentrations. Conversely, the metabolism of phenylalanine, umbelliferone, p-coumaric acid, naringenin, resveratrol, and Thn C was more active in accessions containing PD. Correlation analysis between PD and these metabolites offered valuable insights into their interrelationships. Transcriptome analysis revealed specific genes involved in phenylpropanoid biosynthesis, flavonoid biosynthesis, and stilbene biosynthesis pathways that are crucial for PD biosynthesis. Moreover, the identification of cyclase and ketoreductase genes, which were up regulated in accessions with high PD, provided further understanding of the biosynthetic pathway leading to PD. Finally, based on metabolite levels and gene expression data, a hypothetical pathway from resveratrol to Thn C was proposed, suggesting its potential involvement in PD biosynthesis.

**Highlight:** The metabolic and transcriptomic analysis led to the identification of specific metabolites and genes that are involved in the biosynthesis of dihydrisocoumarin.

## Introduction

The sessile nature of plants has led to the evolution of diverse pathways that have enabled them to respond to environmental stimuli and establish sophisticated relationships with co-evolving species through the production of specialized biomolecules not generally involved in primary metabolism (Dixon, 2001; Hartmann, 2007; Weng *et al*, 2021). These secondary metabolites include an array of more than 200,000 diverse chemical compounds derived from multiple biosynthetic pathways (Dixon, 2001; Schwab, 2003). However, the origin of most of these biomolecules is attributed to phenylpropanoid biosynthesis. Phenylpropanoids are a group of organic compounds derived from the amino acid L-phenylalanine through a deamination process facilitated by L-phenylalanine ammonia lyase (PAL). This non-oxidative deamination of phenylalanine converts phenylalanine to trans-cinnamate and directs carbon flow to various branches of the general phenylpropanoid pathway (PPP) (Fig 1, Dixon *et al*, 2002; Vogt, 2010). Thus, the PPP acts as a central hub, channeling primary metabolites through various enzyme-mediated bioconversions to form specialized metabolites, priming the plant to tackle various biological hurdles. A large number of these bioactive molecules has therapeutic properties. This includes flavonoids, stilbenes and polyketides such as 3,4-dihydroisocoumarins and melleins (Dixon & Piva, 1995; Austin & Noel, 2003; Noel *et al*, 2005; Dewick, 2009; Noor *et al*, 2020). An important class of compounds derived from the PPP pathway are the coumarins, which belong to the polyphenol class of secondary metabolites (Bourgaud *et al*, 2006). In the plant kingdom, coumarins have been reported in approximately 150 different species from nearly 30 different families, including Apiaceae, Caprifoliaceae, Clusiaceae, Guttiferae, Nyctaginaceae, Oleaceae, Rutaceae and Umbelliferae. The distribution of these bioactive molecules in plant tissues, whether it’s leaves, seeds, fruits, roots, or latex, depends on the specific type of coumarin, the plant species, and its developmental stage (Venugopala *et al*, 2013). Extensive research has been conducted to gain insight into the ability of coumarins to facilitate iron acquisition from soils in Nicotiana (Kai *et al*, 2006; Rajniak *et al*, 2018; Lefèvre *et al*, 2018), modify the rhizosphere to create a favourable environment for certain microbes in Arabidopsis (Voges *et al*, 2019), confer resistance to fungal infections in Brassica sp. (Tortosa *et al*, 2018), and scavenge reactive oxygen species under abiotic stress conditions (Fourcroy *et al*, 2014; Döll *et al*, 2018). While the plant kingdom remains an important source of chemical compounds, recent studies have highlighted the utility of natural coumarins and their synthetic analogs in various fields, including pharmacology, food industry, perfumery, and scientific research (Floc’h *et al*, 2002; Lončar *et al*, 2020; Sugiyama *et al*, 2023; Onder *et al*, 2023). Among the coumarins, isocoumarins represent a class of compounds that are isomeric to coumarins and are characterized by an inverted lactone moiety. While coumarins typically originate from the phenylpropanoid pathway, such as umbelliferone, isocoumarins have their roots from polyketide biosynthetic pathway (Fig. 1, Wu *et al*, 2016; Song *et al*, 2017). Among this, phyllodulcin (PD) is a dihydroisocoumarin present uniquely and abundantly in *H. macrophylla*. In Japan, a traditional sweet herbal tea called amacha containing PD is brewed from dried leaves of *H. macrophylla*. This tea is known for its distinctive sweet taste and associated health benefits (Bassoli *et al*, 2008). In particular, PD is exceptionally sweet, approximately 400-800 times sweeter than sucrose. The sweet taste of the molecule has been attributed to the presence of the orthohydroxymethoxyphenyl (isovanillyl) unit (Bassoli *et al*, 2008; Kim *et al*, 2017; Ciçek, 2020). In *Hydrangea* leaves, PD is naturally present in the form of phyllodulcin-β-D-glucoside. When the plant experiences various stresses such as drought, wounding or senescence, native glucosidases within the plant hydrolyze these glucosides, converting PD to its aglycone form, which has a pleasantly sweet, minty taste. In addition, PD has been shown to have several beneficial properties in both traditional and modern medicine, including antibacterial, antimalarial, antifungal, antiulcer and anti-inflammatory effects (Kawamura *et al*, 2002, Zhang *et al*, 2007, Kim *et al*, 2017, Cho *et al*, 2023).

**Figure 1.**
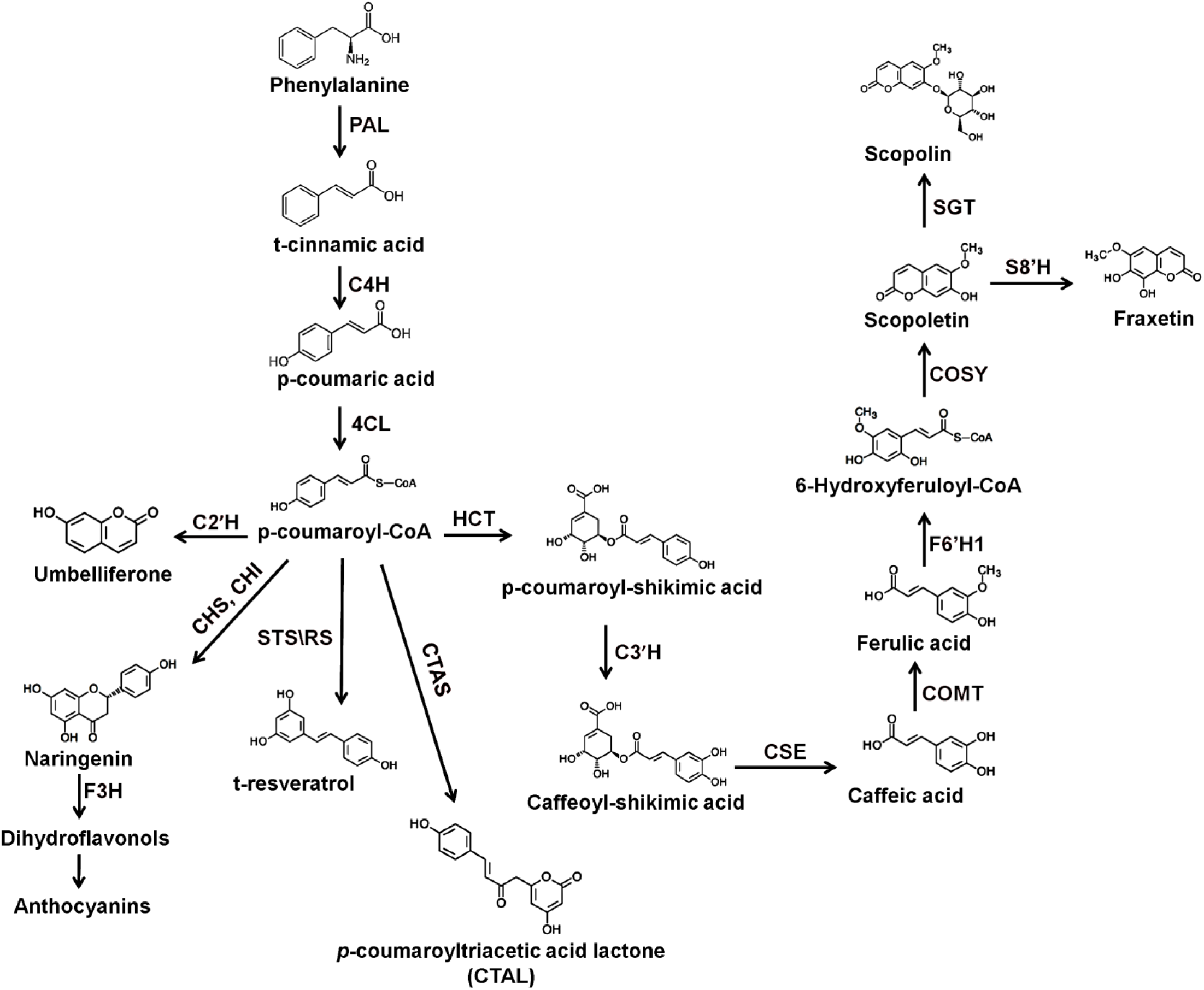
Schematic representation of phenylpropanoid metabolism in plants leading to the formation of coumarins, flavonoids and polyketides. The corresponding enzymes are shown in bold. Phenylalanine is the starting point for the synthesis of different Products, while coumaroyl-CoA is the branching point for different pathways. 4CL, 4-coumarate-CoA ligase; C3H, p-coumaroyl shikimate 3′ hydroxylase; C4H, cinnamic acid 4-hydroxylase; CHS, chalcone synthase; CHI; chalcone isomerase; COMT, caffeate/5-hydroxyferulate 3-O-methyltransferase; CSE, caffeoyl shikimate esterase; F3H, flavanone 3-hydroxylase; HCT, hydroxycinnamoyl transferase; PAL, Pheniylalanine ammonia lyase; RS, resveratrol synthase; C2H, cinnamate 2 hydroxylase; F6’H1, feruloyl-CoA 6’-hydroxylase; SGT, scopoletin glucosyl transferase; S8H, scopoletin 8-hydroxylase; CTAS, p-coumaroyltriacetic acid synthase; COSY, coumarin synthase.

Like its parent biomolecule class, PD originates from the PPP. Early studies, aided using labelled ^14^C compounds, suggested that the initiation of PD biosynthesis occurs via L-phenylalanine and cinnamic acid (Basyouni *et al*, 1964; Yagi *et al*, 1977). These studies suggested that branching from p-coumaric acid was a possible route for PD biosynthesis and that hydrangenol (HD) could serve as a precursor in this pathway. This conclusion was drawn based on the incorporation ratio of labelled carbon, which indicated that the introduction of the C-3’ hydroxy group in PD occurs after the formation of HD. It was also reported that three molecules of malonyl-CoA condensed to form a tetraketide intermediate, which was probably involved in HD biosynthesis (Ibrahim and Towers, 1960). In 1999, researchers identified an enzyme called p-coumaroyltriacetic acid synthase (CTAS) in *H. macrophylla*. This enzyme catalyzes three consecutive decarboxylation reactions of p-coumaroyl-CoA and malonyl-CoAs, ultimately leading to the production of p-coumaroyltriacetic acid tetraketide (CTA), which is further converted to p-coumaroyltriacetic acid lactone (CTAL) (Fig. 1, Akiyama *et al*, 1999). Although an enzyme responsible for stilbene carboxylic acid biosynthesis in *Hydrangea* has not been identified, it has been suggested that CTAS, in association with a hypothetical polyketide cyclase (PKC) and a ketoreductase (KR), may be involved in hydrangeic acid formation (Akiyama *et al*, 1999; Austin and Noel, 2003). A reduction step was also suggested by researchers involving the enzyme stilbene carboxylate synthase (STCS) which converts dihydro-p-coumaric acid to stilbene carboxylates such as 5-hydroxy-lunularic acid (Eckermann *et al*., 2003). Hydrangeic acid could potentially lead to the formation of HD and subsequently PD, as PD can be derived from HD by C-3’-hydroxylation and C-4’O-methylation (Yagi *et al*., 1977). In parallel, thunberginols synthesized from resveratrol have also been proposed to be involved in PD biosynthesis (Çiçek *et al*, 2018; Preusche *et al*, 2022). While these studies lay the groundwork for unravelling the PD biosynthetic pathway in *Hydrangea*, a comprehensive understanding requires further research into the genes and potential intermediates involved in this pathway. For these reasons, the following study was undertaken to understand the dynamics of PD and HD metabolism in *Hydrangea* plants, which are of both scientific and industrial importance. In addition to optimizing harvest time for efficient utilization and processing methods, we identified potential genes and associated metabolites involved in the PD biosynthetic pathway.

## Materials and Methods

### Plant material and sample processing

Fresh, fully expanded young upper leaves from 182 *Hydrangea macrophylla* accessions were obtained from Koetterheinrich Hortensienkulturen (Lengerich, Germany) and subsequently screened for PD and HD concentrations after drying the leaf tissue. Thirteen accessions were selected and were chosen for metabolomic and transcriptomic experiments (Tables S3a and S3b). From the selected 13 accessions, five accessions of *H. macrophylla*, namely VAR-552, VAR-746, VAR-553, VAR-212 and VAR-163, were grown from the cutting stage in the greenhouse at IPK-Gatersleben under controlled conditions, with an 18 h photoperiod and a light intensity of 200 μmol m^-2^s^-1^, a temperature of 21/19°C (day/night) and a relative humidity of 60%. The plants were randomized and grouped daily. The plants were allowed to grow for 75 days, at which point freshly expanded leaves were harvested, immediately frozen, ground to a fine powder and stored at −80°C until use. These materials were used for metabolomic and transcriptomic analysis.

### Chemicals

LC-MS grade acetonitrile, methanol, and n-hexane used in this experiment were procured from Carl Roth (Karlsruhe, Germany). Formic acid was obtained from Thermo Fisher Scientific (Germany). Analytical standards essential for the quantitative measurements of PD and HD were provided by Symrise AG, (Dr. Ley, Holzminden, Germany). Additionally, the analytical standards required for phenylpropanoid analysis were purchased from Sigma Aldrich (Merck AG, Taufkirchen, Germany).

### Extraction of PD and HD

Both PD and HD were extracted with slight modifications to an existing method that included drying and fermentation steps, specifically using accelerated solvent extraction (ASE) as described by Lee *et al*. (2007) and Jung *et al*. (2016). Briefly, 5-10 mg of powdered tissue, both fresh and dried, was fermented with 0.2 mL of ultrapure water for 2 h at 40°C. Then, 1.8 mL of methanol was added and the mixture was incubated in an ultrasonic bath for a further 2 h at 40°C. The supernatant was separated by centrifugation at 13,000 RPM for 15 minutes and 1 mL of methanol was added to the sediment, which was then sonicated for 1 h. The supernatant from this fraction was combined with the previous fraction after centrifugation at 13,000 RPM for 15 minutes. The final mixture was collected and passed through Strata C18 columns (55µm, 70Å, 100 mg/ml, Phenomenex, Germany) preconditioned with 1 mL of methanol and eluted with 1 mL of methanol. The final volume was collected and subjected to LC-MS analysis.

### Extraction of phenolic compounds

For the extraction of phenolics, including p-coumaric acid, trans-cinnamic acid, caffeic acid, ferulic acid, naringenin and trans-resveratrol, a liquid extraction with methanol was performed with slight modifications based on the method described by Irakli *et al*. (2021). 5-10 mg of finely ground plant tissue was combined with 1 mL of 80% methanol and stirred in an ultrasonic bath for 1 h at 30°C. The resulting extract was then centrifuged at 13,000 RPM for 15 min at 4°C and the extraction process was repeated. The supernatant was then filtered through a 0.45 µm membrane filter and collected in new Eppendorf tubes for further analysis.

### Extraction of coumarins

Extraction of scopolin, scopoletin, esculin, esculetin, fraxetin and umbelliferone was performed with slight modifications based on a previously published method (Perkowska *et al*, 2021). Briefly, 1 mL of methanol was added to the finely ground and powdered samples, which were then subjected to sonication for 1 h, followed by incubation in the dark at 4°C for a further 2 h. All samples were then centrifuged at 13,000 RPM for 15 min, and the resulting supernatants were carefully transferred to new Eppendorf tubes. For further processing, the extracts were dried in a vacuum centrifuge for 2 h at 45°C (Christ, RVC 2-33 RI, Germany). Subsequently, 100 μL of 80% methanol was added to the dried extracts to dissolve the compounds and incubated overnight at 4°C. The next day, the extracts were vortexed for 10 minutes and separated into 50 μL aliquots. These samples were stored at −20°C until analysis by LC-MS.

### UPLC-MSMS analysis

UPLC-MSMS analyses were performed on an Agilent 1290 UPLC system coupled to an Agilent 6490 triple quadrupole mass spectrometer. The chromatographic separation was performed on an ZORBAX RRHD Eclipse Plus C18, 95Å, 2.1×50 mm, 1.8 µm column at a flow rate of 0.45 mL/min and a column temperature of 40°C (Agilent Technologies, Waldbronn, Germany). The separation was performed with a gradient of solvent A (water) and B (acetonitrile), both containing 0.1% formic acid (v/v). The initial percentage of B was 10%, increased linearly to 80% in 5 min and then re-equilibrated to the original conditions for 6 min. ESI-MS/MS analysis was performed in positive and negative ionization mode using nitrogen as drying and nebulizing gas. The gas flow was set at 12.0 l/min at 250°C and the nebulizer pressure was 30 psi. The capillary voltage was 2 kV and the residence time was 20. MassHunter optimizer software was used to select precursor ions using MS2 Selected Ion Monitoring (SIM), product ions using product ion scan for each precursor ion and optimum collision energy for each transition using multiple reaction monitoring (MRM) acquisition mode. Product ions were selected as the most abundant ions in a composite product ion scan spectrum obtained for a given precursor ion at multiple collision energies. Five different concentrations of selected 14 metabolites were used to prepare a calibration curve from a range of 0.01-50 µg per ml and the absolute quantification was performed using a single multiple reaction monitoring (MRM) transition for each analyte (Table S5). The limit of quantification (LOQ) for the coumarins were measured by triplicate injections of the standard solutions based on signal to noise ratio of 10. Agilent MassHunter software (B.07.01, Agilent Technologies, United States) was used for data acquisition as well as final qualitative and quantitative analysis.

### RNA isolation, sequencing and analysis

To isolate total RNA from the plant tissues, the Spectrum™ Plant Total RNA Kit, obtained from Sigma Aldrich, was employed following the manufacturer’s guidelines, and it was highly effective for isolating RNA from *Hydrangea* leaf tissues. The complete detailed procedure is described in supplementary protocol S1.

### Validation of gene expression using qRT-PCR

The RevertAid First Strand cDNA Synthesis Kit from Thermo Scientific was utilized to perform cDNA synthesis, and 0.3 µg of total RNA was employed as the starting material. The synthesis was initiated using oligo(dT) primers. The primers for the quantitative real-time polymerase chain reaction (qRT-PCR) were created through primer design software Primer3, and they were subsequently synthesized by the company Metabion (Germany). The list of primers designed for validation with qRT-PCR are represented in Table S2. The detailed procedure is described in Supplementary protocol S2.

### Weighted gene co-expression network analysis (WGCNA)

WGCNA is a data-driven method that discovers co-clustered gene sets (modules) based on weighted correlations between gene transcripts. For the construction of gene co expression networks, weighted gene co-expression network analysis (WGCNA) was performed on 22,980 genes having high variance among the accessions which were obtained from RNA sequencing of *Hydrangea* accessions. The WGCNA (version 1.72-1) R software package is a comprehensive collection of R functions for performing various aspects of weighted correlation network analysis. This R package included functions for network construction, module detection, gene selection, calculations of topological properties, data simulation, visualization, and interfacing with external software (Langfelder and Horvath, 2012).

### Statistical analysis

Statistical analysis was performed using GraphPad Prism 9.5.1 (GraphPad Software Inc.) and R (version 4.0.3). One-way ANOVA with post hoc Tukey’s test (p ≤ 0.05) was used for multiple comparisons. Paired sample t-tests were used to assess the significance of differences between the control and treatment groups.

## Results

### Selection of suitable plants with varying levels of PD and analysis of PD and HD in both fresh and dried leaves

To investigate the distribution of PD and HD, an initial screening was done on the fresh and dried leaves (dried at 40°C for 48 h) of 182 accessions of *H. macrophylla* (provided by Koetterheinrich Hortensienkulturen, Lengerich, Germany, Table S3a and 3b). For further experiments, 13 accessions with different PD and HD levels were selected (Fig. 2, Table 3a and 3b) and *H. paniculata* where PD and HD are absent was used as a negative control. LC-MS analysis was then performed to quantify both PD and HD concentrations in all desired accessions. PD concentrations among the different accessions, measured in the leaves of 75-day-old plants, ranged from 1.36 to 38.34 mg.g^-1^DW, while HD concentrations ranged from 1.28 to 28.80 mg.g^-1^DW (Fig. 2A). Based on these measurements, the selected thirteen accessions were grouped into four different classes according to their PD and HD concentrations: high PD (VAR-552, VAR 746 and VAR-753), high PD/HD (VAR-553, VAR-547, VAR-897 and VAR-908), high HD (VAR-751, VAR-827 and VAR-760) and low PD/HD (VAR-212, VAR-910 and VAR-163).

**Figure 2.**
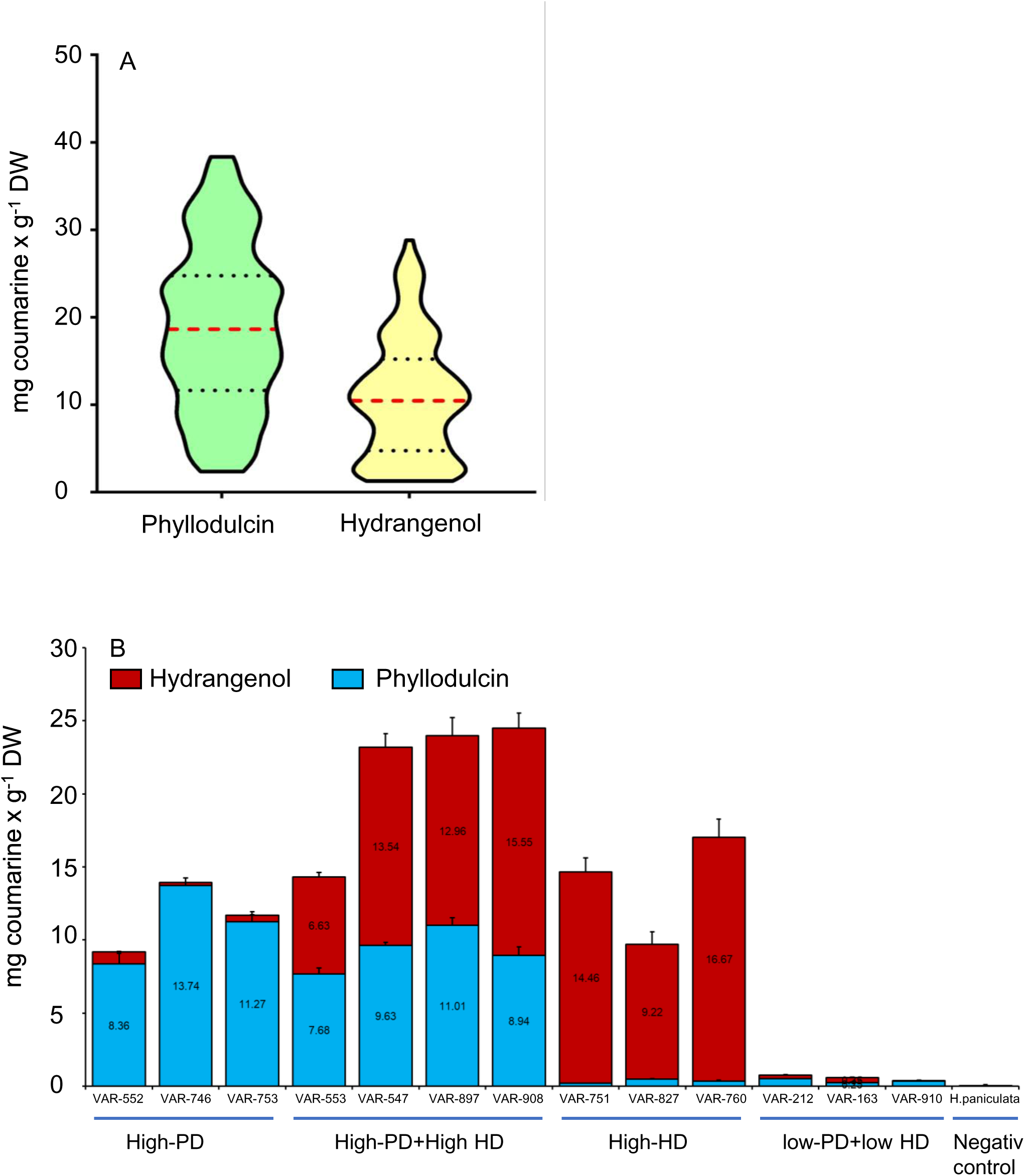
Evaluation and selection of 182 different Hydrangea accessions determined by levels of phyllodulcin (PD) and hydrangenol (HD). (A) In the violin plot, the median observations are depicted by a red dotted line, while the 3rd and 1st quartiles are represented by black dotted lines. (B) Selection of 14 accessions containing different level of PD and HD for further experiments. Blue bars show the content of phyllodulcin while red bars show the content od HD. H.paniculata was serving as the negative control.

A comparison of freshly harvested leaves with dried leaves revealed that the dried leaves of the selected accessions contained significantly higher levels of PD and HD, up to 8-fold (Fig. S1A, B), compared to their fresh counterparts (Fig. S1C, D). Notably, PD and HD were not detected in *H. paniculata* (Fig. S1A-D). The screening and selection process of *Hydrangea* accessions served to optimize the research workflow and acted as a valuable resource for the development of next generation crosses with increased PD and HD content. It also provided insight into the enhancement of PD and HD content by leaf desiccation.

### Distribution of phenylpropanoids in *Hydrangea* accessions

To investigate the changes in phenylpropanoid metabolism in selected *H. macrophylla* accessions, the concentrations of 14 closely related phenylpropanoids were analyzed in freshly harvested leaves. In particular, the high PD, high HD and high PD/HD accessions showed significantly increased concentrations of phenylalanine, which serves as a starting compound in phenylpropanoid metabolism (Fig. 3A). Furthermore, all low PD/HD accessions and *H. paniculata* accumulated significantly higher concentrations of trans-cinnamic acid compared to the other groups (Fig. 3B). In plants, activated p-coumaric acid is branched to various metabolites, including umbelliferone, caffeic acid, naringenin chalcone and resveratrol. It was evident that plants with high PD and/or HD concentrations contained significantly higher levels of p-coumaric acid and naringenin compared to low PD/HD accessions and *H. paniculata* (Fig. 3C, D). The concentrations of scopoletin and fraxetin were higher in low PD/HD accessions and *H. paniculata* compared to high PD accessions (Fig. 3E, F). The levels of caffeic acid and ferulic acid were significantly higher in low PD/HD accessions compared to all other accessions (Fig. 3G, H). The levels of stilbenoids, especially resveratrol, were lower in the low PD/HD accessions compared to all other plants and were not detected in *H. paniculata* (Fig. 3I). Among the five different thunberginols, Thn C showed clear differences between different *Hydrangea* accessions. The relative abundance of Thn C was lower in low PD/HD accessions compared to all other plants and was not detected in *H. paniculata* (Fig. 3J). In addition, high PD and/or HD accessions showed significantly higher levels of umbelliferone (Fig. S7A), whereas the concentration of scopolin was higher in low PD/HD accessions and *H. paniculata* compared to high PD accessions (Fig. S7B). Accessions with high PD showed higher levels of esculin and lower levels of esculetin compared to other study groups (Fig. S7C, D)

**Figure 3.**
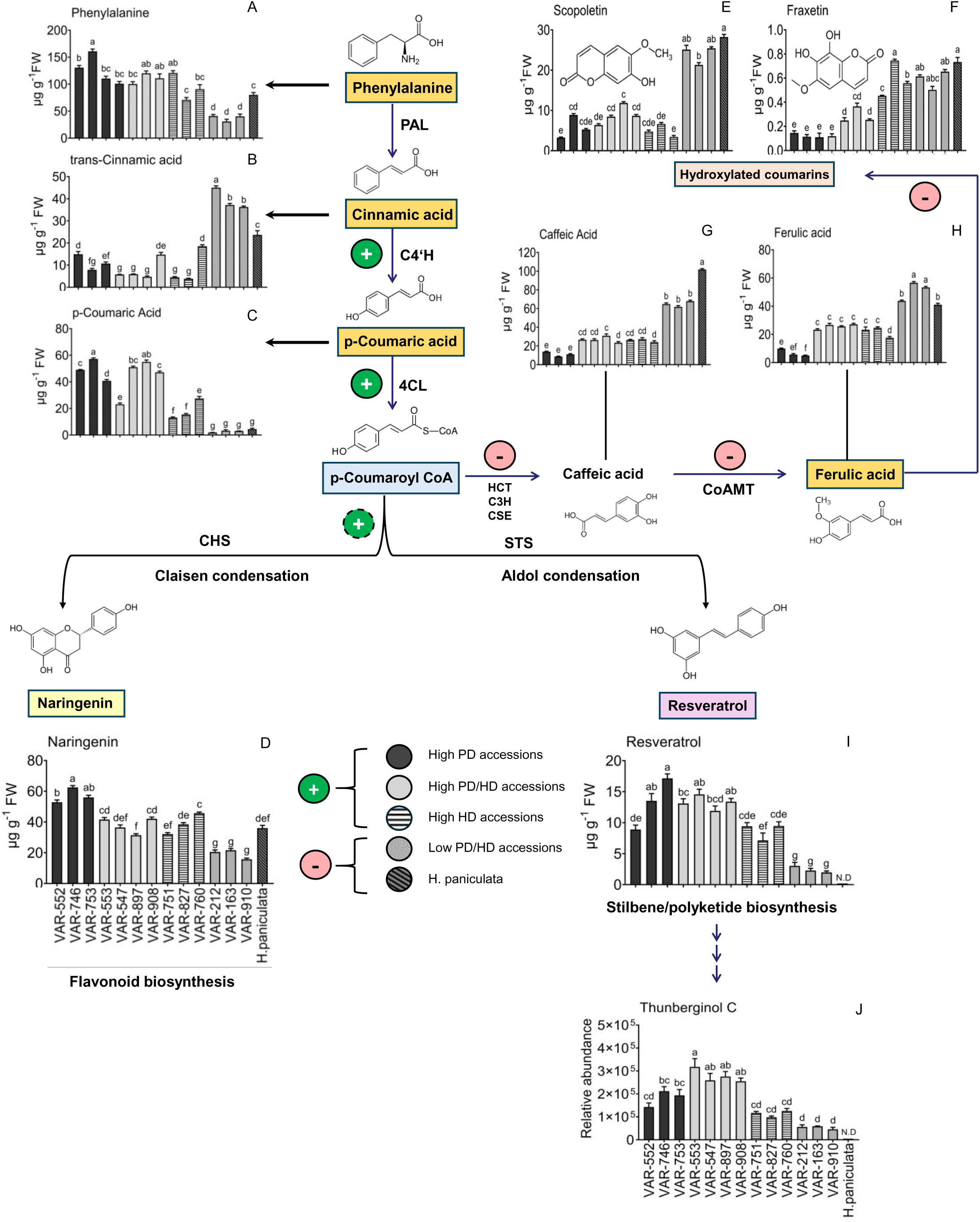
Levels of specific phenylpropanoids, determined in different Hydrangea accessions. The concentrations of (A) phenylalanine, (B) transcinnamic acid, (C) p-coumaric D) naringenin, (E) scopoletin, (F) fraxetin, (G) caffeic acid, (H) ferulic acid, (I) trol and (J) thunberginol C measured in 14 selected Hydrangea accessions. Analysis rformed on freshly harvested, fully expanded, young upper leaves. Bars represent the f 6 independent biological replicates (n=6) and standard error. Not detected (N.D.) indicates the absence of the corresponding metabolite. Dark bars show accessions with high PD, light grey bars accessions with high PD/HD, hatched bars accessions with high HD, dark gray bars accessions with low PD/HD and dark hatched bars H. paniculate as negative control. Different letters indicate significant differences between accessions according to one-way ANOVA followed by post-hoc Tukey’s test (p<0.05).

These changes observed in the metabolite profiles of different *H. macrophylla* accessions helped predicting the direction of metabolic conversions within the major PPP, ultimately leading to the biosynthesis of PD and HD.

### Relationship between PD and other metabolites in *Hydrangea* accessions

To assess the interdependence of different phenylpropanoid intermediates and to identify potential contributors to the biosynthesis of PD, a correlation analysis was performed using the metabolite concentrations of different *Hydrangea* accessions (Fig. 4A). In particular, low PD/HD accessions had higher concentrations of cinnamic acid, caffeic acid, ferulic acid, scopoletin, scopolin, fraxetin and esculetin compared to all other accessions, while high PD accessions had higher concentrations of umbelliferone, naringenin and resveratrol (Fig. 4A).

**Figure 4.**
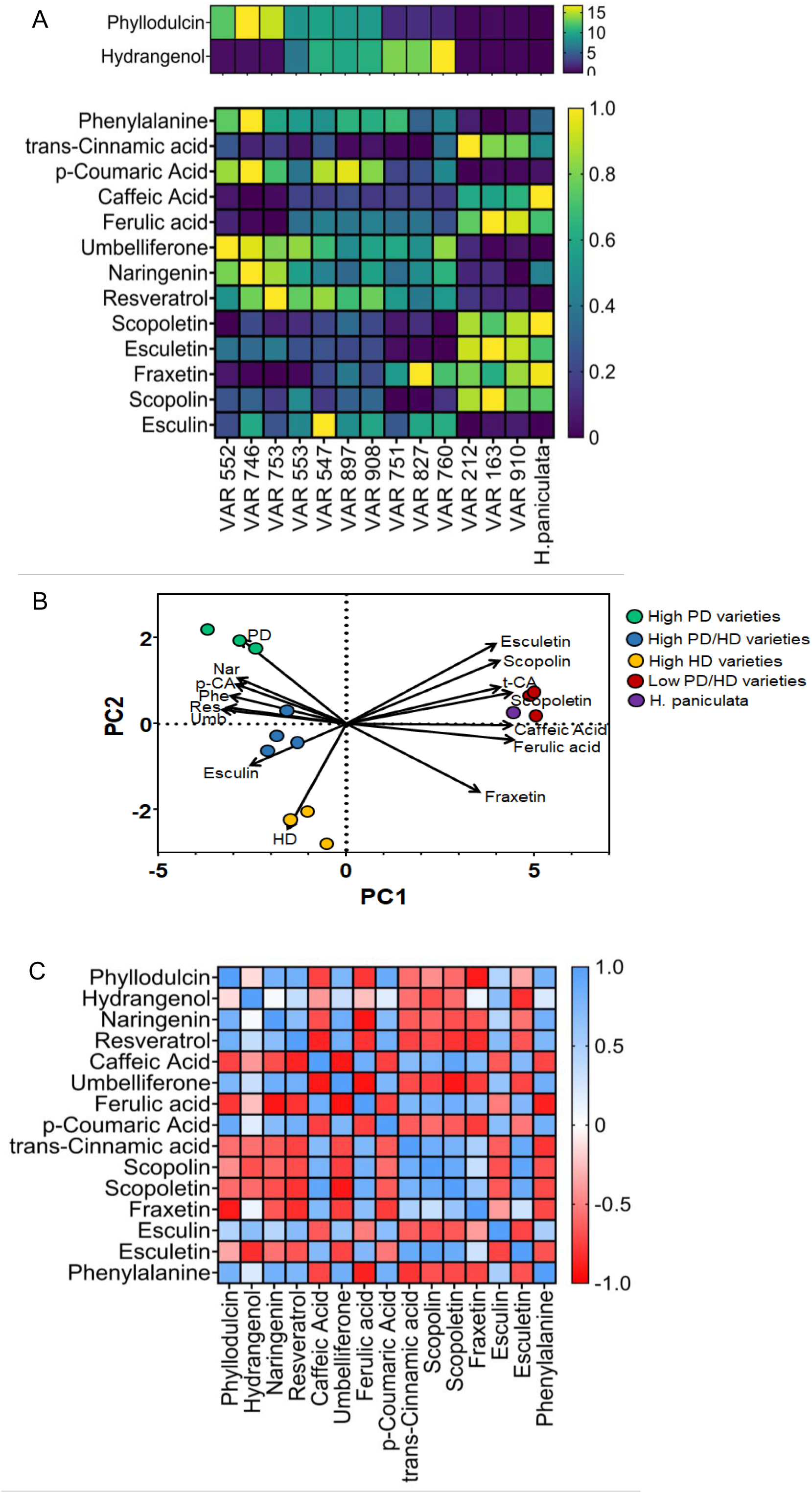
(A) Variation in phenylpropanoid derivatives among Hydrangea accessions. Heat map of absolute concentrations of PD and HD (upper panel) and heat map of relative concentrations of 13 metabolites in different Hydrangea accessions lower panel). Yellow colour represents the highest relative concentration and dark purple represents the lowest relative concentration of metabolites. (B) PCA biplot showing different Hydrangea accessions as PCA plot and metabolite concentrations as loading Each point represents different Hydrangea accessions and the arrows represent eigenvectors of 15 metabolites. The abbreviations Nar, p-CA, Phe, Res, Umb and t-CA represent naringenin, p-coumaric acid, phenylalanine, resveratrol, umbelliferone and trans-cinnamic acid, respectively. (C) Correlation analysis between phenylpropanoids in Hydrangea accessions. Pearson’s correlation analysis was performed between 15 metabolite concentrations of different Hydrangea accessions. Blue colour represents high positive correlation (r=1) and red colour represents high negative correlation (r=-Metabolite analysis was performed on freshly harvested fully expanded upper leaves (n=6).

Principal component analysis (PCA) was performed using the concentrations of all selected metabolites in each accession to group accessions based on their biochemical diversity. Principal component 1 (PC1) captured 70.25% of the variance in metabolite concentrations between accessions and principal component 2 (PC2) captured 15.48% of the variance in different Hydrangea accessions. The PCA plot showed that high PD, high PD/HD, low PD/HD and high HD accessions formed distinct clusters, indicating that metabolite concentrations were a major factor contributing to the differences between these groups (Fig. 4B). In the loading plot, the vector angles for PD and HD were nearly orthogonal to each other, indicating a lack of correlation between their concentrations among the accessions. Accessions VAR-552, VAR-746 and VAR-753 clustered along the PD vector projection, suggesting that PD may be a key factor for this grouping (Fig. 4B). Similarly, VAR-751, VAR-760 and VAR-827 clustered closer to the HD eigenvector projection, suggesting that HD may be responsible for this clustering. Accessions VAR-212, VAR-163, VAR-910 and *H. paniculata* clustered close to the trans-cinnamic acid, scopolin and caffeic acid eigenvectors. On the other hand, VAR-553, VAR-908, VAR-897 and VAR-547 clustered closer to the eigenvectors of esculin, umbelliferone, resveratrol and phenylalanine (Fig. 4B).

Furthermore, the correlation analysis between the compounds involved in the PPP showed strong and positive correlations between PD concentration and the concentrations of naringenin, resveratrol, umbelliferone, p-coumaric acid, esculin and phenylalanine (Fig. 4C). Conversely, there were strong negative correlations between caffeic acid, ferulic acid, trans-cinnamic acid, scopolin, scopoletin, fraxetin and esculetin with PD concentration (Fig. 4C).

### Differential expression of genes involved in PPP and associated pathways

To investigate the genetic regulation of the PPP and related pathways in different *Hydrangea* accessions sorted by their biochemical content, a clustering heat map analysis was performed on the transcriptome data with all accessions. Gene clustering revealed distinct gene expression patterns among the different accessions. In particular, *H. paniculata*, which lacked PD showed significantly different gene expression with strong down-regulation of genes (green spots) compared to other accessions containing high or significant amounts of PD, while in all other accessions different clusters of up and down regulated genes were observed, independent of the amount of PD (Fig. 5A).

**Figure 5.**
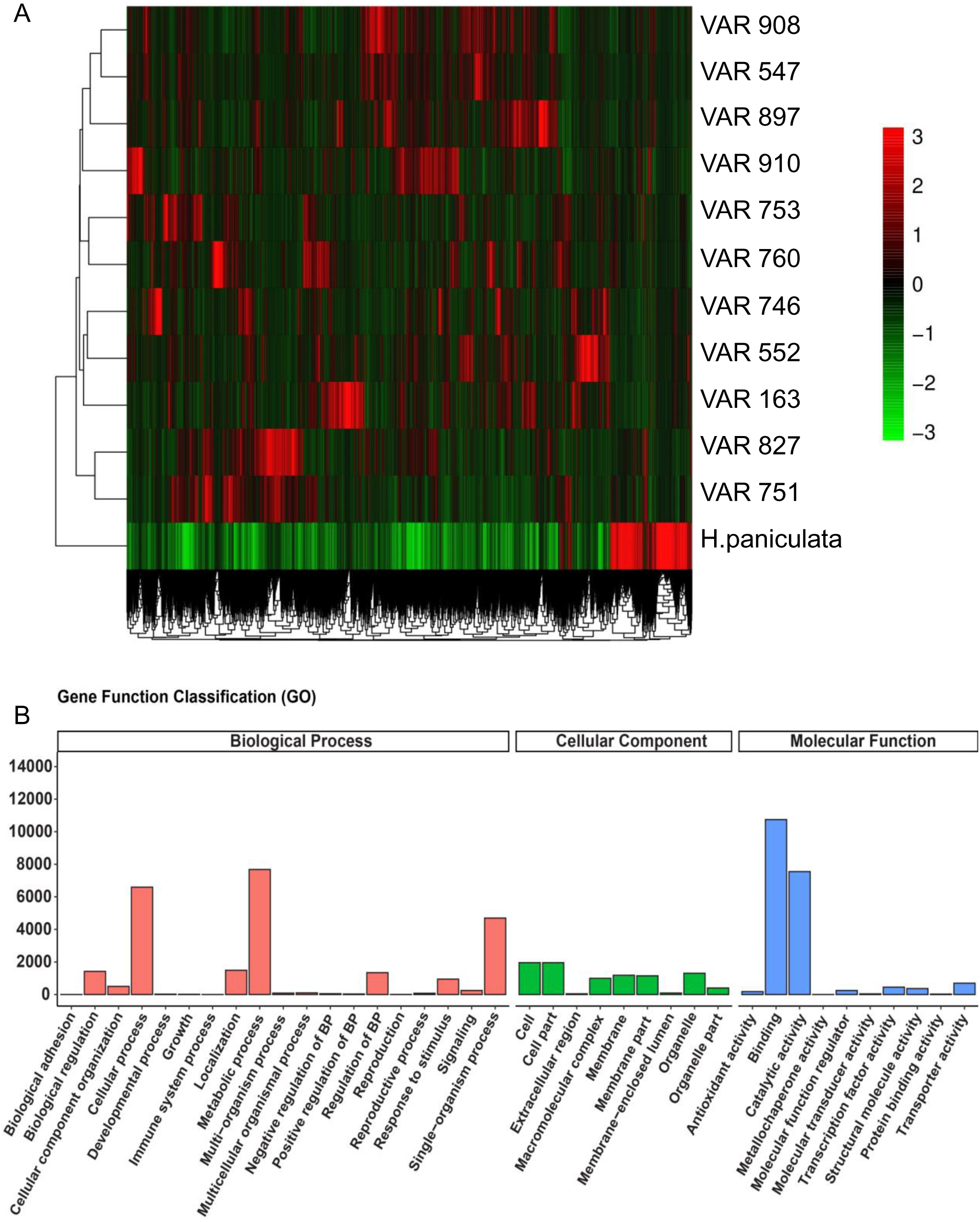
Differences in gene co-expression patterns among Hydrangea accessions. (A) Heatmap showing the relative gene expression patterns in H. paniculata and 11 different accessions of H. macrophylla. Normalized expression data using fpkm values of genes were used to perform the up- and down regulated genes. Green colour represents downregulated and red colour represents upregulated genes (B) Gene Ontology (GO) annotation of differentially expressed genes classified by biological process, cellular component and molecular function. The 38 most significant terms were selected for display. The data were obtained from the full length transcriptome of leaf tissues of all Hydrangea accessions. Only GO terms with padj < 0.05 are shown.

The Gene Ontology function classification categorised the genes obtained from RNA sequencing into three main classes: biological process, cellular component and molecular function. The majority of genes were associated with biological processes and molecular functions (Fig. 5B). Among the biological processes, metabolic and cellular processes were highly represented. In the molecular function category, genes related to binding and catalytic activity were predominant.

In addition, KEGG enrichment analysis was performed to assign differentially expressed genes to different biochemical pathways. In most comparisons, pathways related to ribosome, phenylpropanoid biosynthesis, flavonoid biosynthesis, flavone and flavonol biosynthesis, and stilbenoid, diarylheptanoid and gingerol biosynthesis were among the top 20 most highly expressed metabolic pathways with corrected p-value (padj) < 0.05 (Fig. S2, red large dots). A similar phenomenon was observed when comparing the high PD/HD or high HD accessions with the very low PD accessions. In contrast, these pathways were not significantly enriched when comparing the transcriptome profiles of accessions containing PD and/or HD with the negative control *H. paniculata* (Fig. S3).

These results indicate that differentially expressed genes associated with flavonoid biosynthesis, phenylpropanoid biosynthesis and stilbene biosynthesis are highly enriched when comparing the transcriptome of a high PD and/or HD accession with that of a low PD and/or HD accession.

### Relationship between genes and metabolites involved in PPP and related pathways

To investigate the relationship between genes involved in PPP and related pathways with metabolite concentrations, Weighted Gene Correlation Network Analysis (WGCNA) was performed, followed by correlation of eigengenes with metabolite levels. WGCNA is an unsupervised analysis method that clusters genes based on their expression patterns and is commonly used to study biological networks through pairwise correlations between variables. To increase the stability of the network, genes expressed as 0 in all samples were removed and high-quality genes with a variance cut-off greater than 0.55 were selected, resulting in 22,980 genes with high variance between accessions (Table S1). A gene cluster dendrogram tree was constructed, with each branch representing a gene cluster with highly correlated expression levels (Figure 6A). The closeness of the branches indicated the similarity between gene sets, and genes with similar expression patterns were grouped together in the same module. In total, 85 modules of different colours were obtained, each containing co-expressing genes. Based on the module eigengene similarity, 16 final expression modules were observed, with the dark olive-green module containing the most genes (5065) and the grey module containing the least (68) (Figure 6B). Pearson’s correlation was performed to estimate module-metabolite relationships.

**Figure 6.**
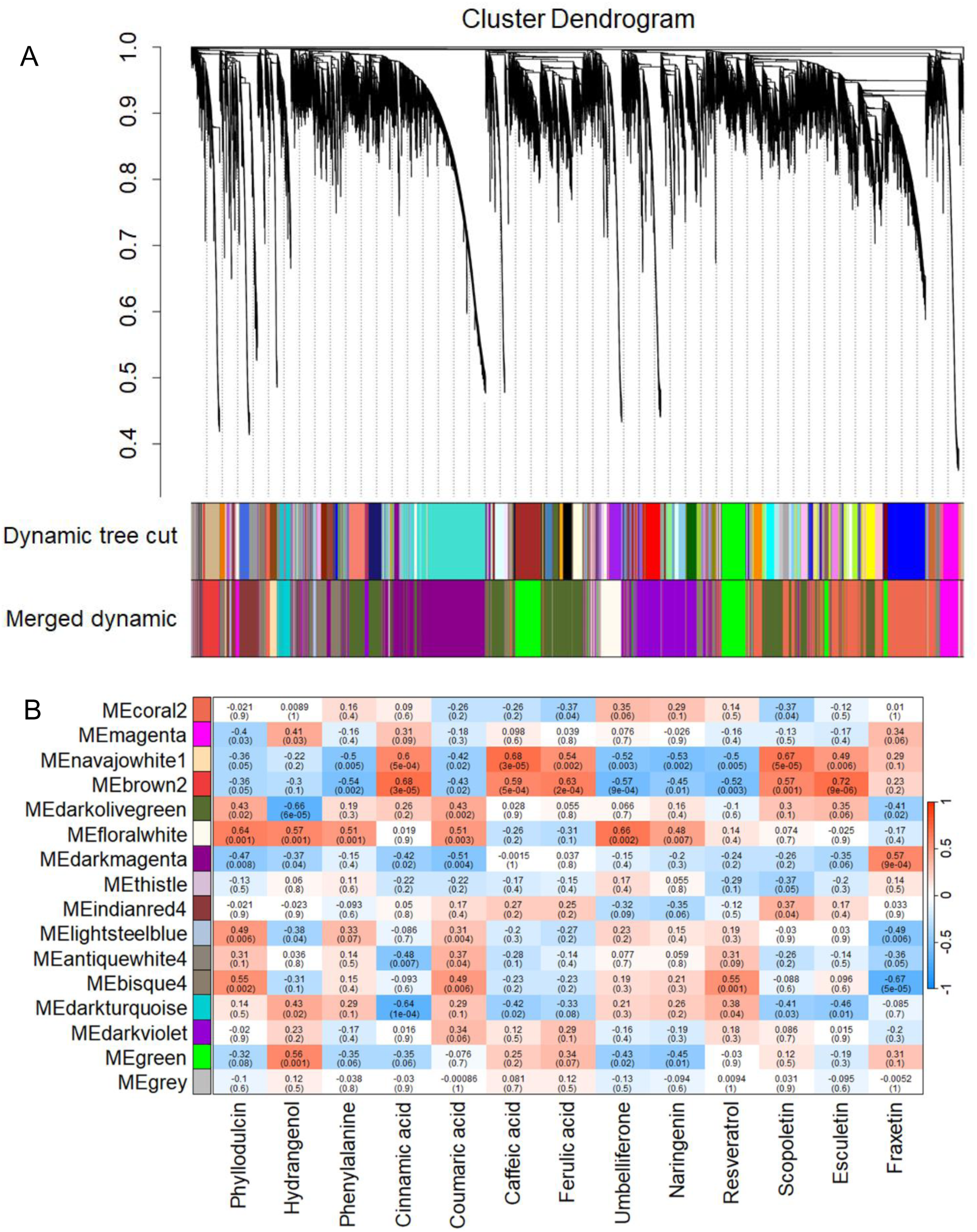
Correlation analysis between the identified genes and metabolites using weighted Gene Co-expression Network Analysis (WGCNA). (A) Hierarchical clustering dendogram of the genes. Gene clustering tree (dendrogram) obtained by hierarchical clustering of adjacency-based dissimilarity to detect 85 co-expression clusters, with corresponding color assignments shown as a dynamic tree section. (B) Modules with strongly correlated eigengenes were merged based on threshold to assign highly co-expressed genes into 16 separate modules. Color bars reflect module assignments before and after the merging of closed modules. Each color represents a module and the gray module indicates none co-expression among the genes. Analysis was done on 22,980 genes obtained from RNA sequencing of 11 H. macrophylla accessions.

Figure 6C and Table S1 show the relationship between the identified genes in different modules including magenta, navajowhite1, brown2, darkolivegreen, floralwhite, darkmagenta, indianred4, lightsteelblue, antiquewhite4, bisque4, darkturquoise and green with the specific metabolite concentrations (p<0.05).

The genes in the floralwhite, bisque4, lightsteelblue and darkolivegreen modules showed a high positive correlation with PD concentration, while the floralwhite, darkturquoise, green and magenta modules correlated with HD concentrations. Phenylalanine concentration positively correlated with the lightsteelblue and floralwhite modules, while p-coumaric acid concentration positively correlated with darkturquoise, floralwhite, lightsteelblue, antiquewhite and bisque4. The navajowhite1 and brown2 modules showed high positive correlation with trans cinnamic acid, caffeic acid, ferulic acid and esculetin concentrations, whereas umbelliferone and naringenin concentrations correlated positively with the floralwhite module. Resveratrol concentration correlated positively with bisque4 and darkturquoise and scopoletin concentration correlated positively with navajowhite1, brown2 and indianred4. Fraxetin concentration correlated positively with the dark magenta module. Notably, the floralwhite module correlated positively with PD, HD, phenylalanine, p-coumaric acid, umbelliferone and naringenin, suggesting a common regulatory network for these metabolites. By analyzing the genes within modules that show a high correlation with each metabolite, we can identify the genes that are involved in the biosynthesis of these metabolites.

### Identification of candidate genes between high and low PD accessions

When comparing the transcriptomes of high PD, high PD/HD, and high HD groups with low PD/HD accessions, the most common and enriched pathways were related to flavonoid biosynthesis, phenylpropanoid biosynthesis, and stilbene biosynthesis. By searching for genes in modules highly correlated with metabolite concentrations and related to these three metabolic pathways, 22 genes were identified (Fig. S8). To understand the regulation of the phenylpropanoid pathway, the expression levels of 16 genes were analyzed in different study groups individually (Fig 7). Among these, the genes involved in stilbene biosynthesis and associated pathways, such as p-coumaroyltriacetic acid synthase (*CTAS*), resveratrol di-O-methyltransferase (*ROMT*), keto reductase (*KR*), type III polyketide synthase (*PKS*) and polyketide cyclase (*PKC*), were upregulated in high PD and/or HD accessions. These genes were of particular interest due to the ability of these enzymes to catalytically produce structurally similar isocoumarins as that of PD and HD.

**Figure 7.**
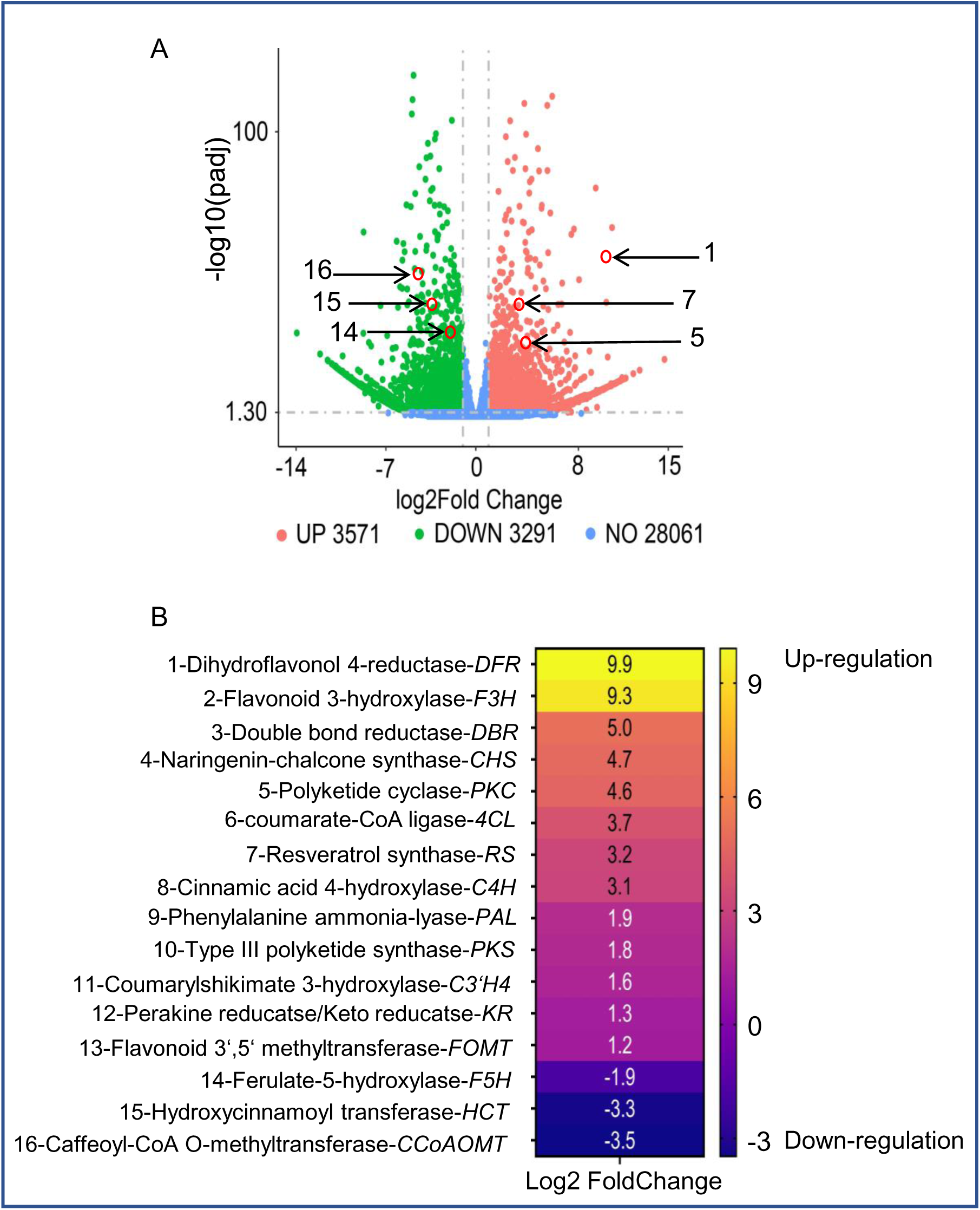
Identification of candidate genes between high and low PD accessions. (A) Volcano plot showing relevant DEGs between the high PD variety VAR-552 and the low PD/HD variety VAR-163. The numbers indicate the genes encoding enzymes in the phenylpropanoid pathway in general and the sub-pathway stilbenoid biosynthesis and flavonoid biosynthesis. Red color shows the up-regulated genes and green color the down-regulated genes. (B) Heat map of candidate genes involved in the phenylpropanoid pathway in the accession with high phyllodulcin content. The numbers and colour code indicate the level of expression, with highest expression in yellow and lowest expression in dark blue. The numbers 1/5/7/14/15/16 indicate the position of the genes in the volcano plot.

### Validation of the expression of candidate genes by quantitative real-time PCR (qRT-PCR)

The genes identified by RNA sequencing were verified by quantitative real-time PCR. For this purpose, specific primers were designed from the sequences of 10 candidate genes involved in phenylpropanoid, flavonoid and stilbenoid biosynthesis, which showed different expression levels in all experimental groups (Table S2). The selected genes were *DFR, DBR, CTAS, PKC, 4CL, PAL1, KR, HCT, CCoAOMT* and *COMT*. For the present study, the most suitable internal reference gene (housekeeping gene) was identified as glyceraldehyde-3-phosphate dehydrogenase (*GADPH*), which has also been reported in previous studies (Zhang *et al*, 2022). Gene expression levels were expressed as fold change and log2 transformed by comparing the same study groups as in the RNA sequencing experiment. Examination of gene expression levels by qRT-PCR revealed that the genes *DBR, CTAS, PKC, 4CL, PAL1* and *KR* were up-regulated when comparing the transcriptome of high PD and/or HD accessions with that of low PD and HD accessions (Fig. S4-S6). Similarly, the genes *HCT, CCoAOMT* and *COMT*, which are involved in caffeic acid synthesis and related downstream metabolites, were found to be downregulated when comparing the same set of accessions (Fig. S4-S6). As an exception, the gene *DFR* was found to be downregulated when comparing accession VAR-746 with VAR-163 (Fig. S4C).

These results show that the expression levels of the transcriptome and RT-qPCR analyses were quite similar to each other, indicating that the genes studied are involved in the metabolic pathways associated with PD biosynthesis.

## Discussion

In the modern era, where food safety is of highest concern, the use of artificial sweeteners in the food industry remains controversial. The recent classification of aspartame, an FDA (Food and Drug Administration)-approved and widely used synthetic sweetener, by the International Agency for Research on Cancer (IARC) in Group 2B as “probably carcinogenic to humans” raises the important question of whether these molecules can continue to be used to enhance the taste of foods (Riboli *et al*, 2023). This scientific curiosity has led researchers to explore a safer and healthier option of mining the plant kingdom for natural sweeteners (Mora and Dando, 2021; Castro-Muñoz *et al*, 2022). An alternative to artificial sweeteners and sugar substitutes is stevioglycoside, a natural sweetener derived from the stevia plant. Stevioglycosides are exceptionally sweet (up to 300 times sweeter than sucrose), low in calories (0.21 kcal/g) and non-cariogenic. However, concerns about the safety of stevioglycosides have led to the establishment of an Acceptable Daily Intake (ADI) of 4 mg/kg by the European Food Safety Authority (EFSA) and the Joint Expert Committee on Food Additives (JECFA) due to potential health effects from the degradation product steviol (Pezzuto *et al*, 1985; Philippe *et al*, 2014). The extraction process for stevioglycosides currently produced in China requires a significant amount of water, involves the use of formaldehyde for preservation (not permitted in Germany) and includes controversial practices such as the use of aluminium salts during precipitation. This process chemically alters the plant material and deviates from the natural composition of *Stevia rebaudiana*. This departure from naturalness could be considered deceptive and potentially in violation of food regulations (DGE/Dokumentations-band_2014.DHCf). A promising alternative is PD, which uses molasses directly from the leaves rather than isolating the active ingredient PD. PD, the sweet component of *H. macrophylla*, is 600 to 800 times sweeter than sucrose and has the potential to be used more widely as a natural sweetener (Yamato *et al*, 1975; Yasuda *et al*, 2004; Kim *et al*, 2017; Preusche *et al*, 2022; Tsukioka and Nakamura, 2023; Cho *et al*, 2023). The anti-diabetic, anti-ulcer and anti-fungal effects and the potential to reduce SARS-CoV-2 infection of PD in *H. macrophylla* extracts and its role in Japanese traditional medicine further justify the importance of studying the biochemical profile and genetic regulation of PD biosynthesis in the plant (Nozawa *et al*, 1981; Zhang *et al*, 2007; Yano *et al*, 2023). So far, the relationship between metabolites and genes in *Hydrangea* sp. has not been addressed together in a single framework, especially by giving priority to PD. Therefore, the work described here was aimed at identifying different accessions with high PD and elucidating the associated biosynthetic pathway.

### Regulation of PD and HD biosynthesis in *Hydrangea* depends on accession-specific expression of genes involved in phenylpropanoid, flavonoid and stilbenoid biosynthesis

Understanding the biosynthesis of PD and HD in Hydrangeas promises to unlock a wide range of bioactive compounds with diverse applications ranging from the food industry to medicine. Previous studies have elucidated the branching of the PPP leading to the production of PD (Fig. 1). Studies using ^14^C-labelled compounds have shown that downstream gallic acid accumulates more labelled carbon, suggesting that the pathway starts with L-phenylalanine and cinnamic acid (Basyouni *et al*, 1964; Yagi *et al*, 1977). These studies also pointed to p-coumaric acid, rather than caffeic acid, which is synthesized from activated p-coumaric acid in plants, as the likely precursor for PD biosynthesis (Fig. 1). This finding is valuable because it suggests that the branch leading to caffeic acid is less favored during PD biosynthesis.

In the current study, a comparative transcriptome analysis between accessions characterized by high PD and/or HD levels and those with low PD/HD levels revealed the significant enrichment of three metabolic pathways: phenylpropanoid biosynthesis, flavonoid biosynthesis and stilbene biosynthesis, as indicated by KEGG enrichment analysis (Fig. S2 and S3). The differences in gene expression between *H. macrophylla* accessions and *H. paniculata* were clearly observed by hierarchical clustering (Fig. 6A). The enrichment of GO terms specific to the metabolic process indicates that the transcripts related to these biological processes are expressed among the accessions (Fig. 5B). Furthermore, the genes of enzymes associated with these pathways showed correlations with the levels of the studied metabolites, as determined by Weighted Gene Co-expression Network Analysis (WGCNA) (Fig. 6). In particular, an increase in phenylalanine levels was observed in accessions with high levels of PD and/or HD (Fig. 3A). In addition, a positive correlation was observed between the concentrations of PD, HD and phenylalanine, confirming the essential role of phenylalanine in PD biosynthesis, with accessions possessing higher phenylalanine producing more PD (Fig. 4B, C). Accessions with high levels of PD and/or HD also showed increased levels of p-coumaric acid (Fig. 3C), and these levels correlated positively with PD and HD levels (Fig. 4B, C). The higher expression levels of *4CL* in high PD and/or HD accessions compared to low PD/HD accessions can be attributed to the activation of p-coumaric acid to p-coumaroyl CoA, which is a critical molecule in the pathway (Fig. 8). These results highlight the significant influence of p-coumaroyl-CoA in PD and HD biosynthesis. Conversely, a negative correlation was observed between PD and HD concentrations and metabolites such as caffeic acid, ferulic acid, scopolin, scopoletin and esculetin, while no correlation was found for fraxetin (Fig. 4). This suggests that these metabolites may have limited or no effect on PD and HD biosynthesis. It is also noteworthy that the concentrations of these biomolecules (except fraxetin) were generally low in accessions with high PD and/or HD levels (Fig. 4).

**Figure 8.**
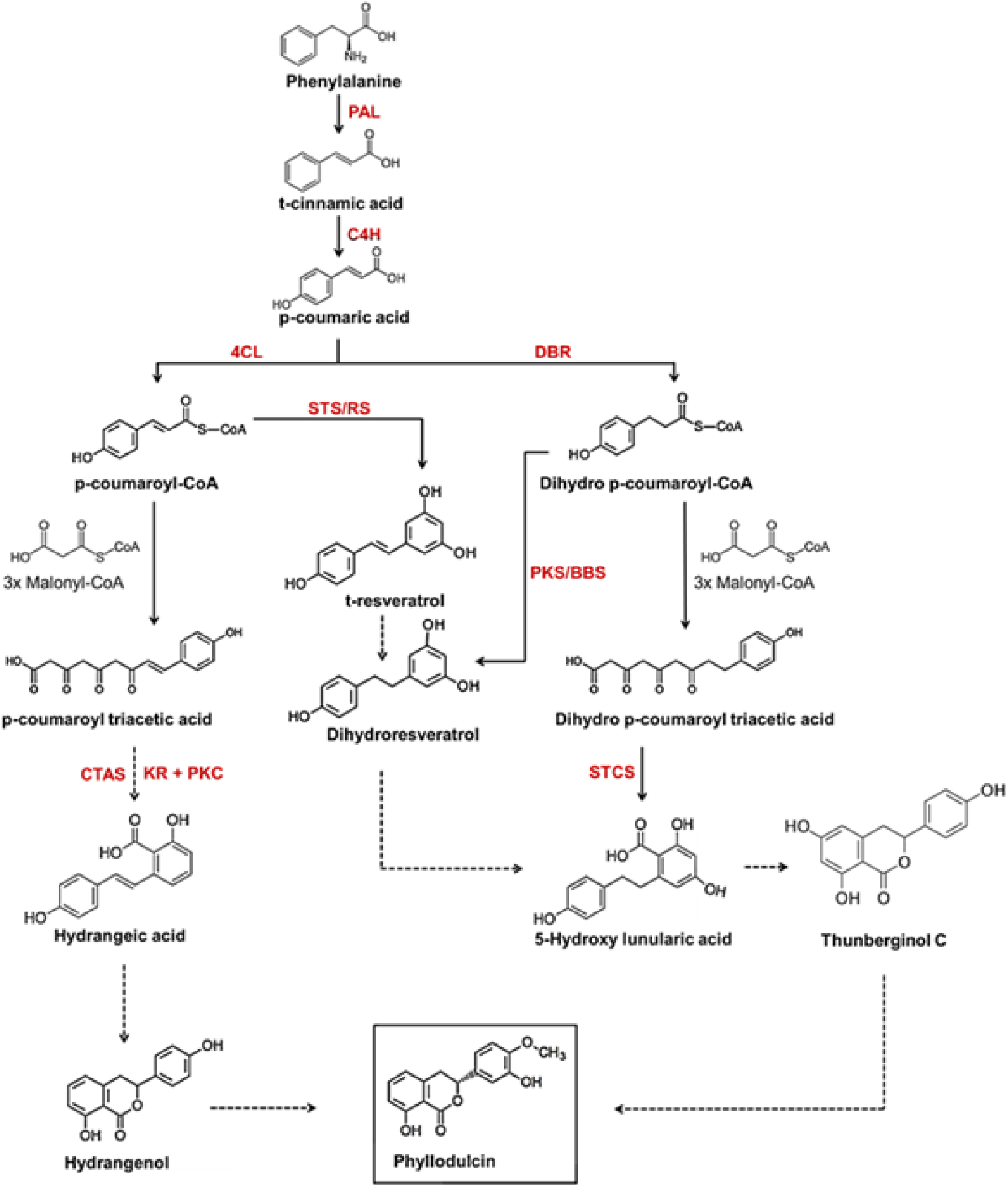
Proposed schematic model of phyllodulcin biosynthesis. Metabolomic and transcriptomic data in the present work led to the following hypothesis. Phenylalanine is converted to p-coumaric acid, the branching point, in three steps catalysed by phenylalanine ammonia lyase (PAL), cinnamate 4-hydroxylase (C4’H) and 4-coumarate-CoA ligase (4CL). P-coumaric acid is either converted to p-coumaroyl-CoA and p-coumaroyl-triacetic acid and finally to hydrangenol, which may serve as a precursor for phyllodulcin formation. P-coumaric acid can also be converted to dihydro-p-coumaroyl-CoA and further to dihydro-p-coumaroyl acid and hydroxylunularic acid or dihydroresveratrol. In addition, p-coumaroyl-CoA can be converted to resveratrol and dihydroresveratrol. The latter and hydroxyunularic acid can be converted to thunberginol C, which is most likely the direct precursor for the synthesis of phyllodulcin. 4CL, 4-coumarate-CoA ligase; STS/RS, stilbene synthases/resveratrol synthase; DBR, double bond reductase; PKS, type III polyketide synthase; BBS, bibenzyl synthase; KR, ketoreductase; PKC, polyketide cyclase; CTAS, p-coumaroyltriacetic acid synthase; STCS, stilbene carboxylate synthase.

At the transcriptome level, the expression levels of *HCT* (p-coumaroyl CoA → p-coumaroyl shikimate, caffeoyl shikimate → caffeoyl CoA), *C3H* (p-coumaroyl shikimate → caffeoyl shikimate), *CCoAOMT* (caffeoyl CoA → feruloyl CoA), *CSE* (caffeoyl shikimate → caffeate), *COMT* (caffeate → ferulate) and *SGT* (scopoletin → scopolin) were found to be downregulated in accessions with high PD and/or HD levels compared to those with low PD/HD levels (Fig. 7A and B). This suggests that in low PD/HD accessions, there is a higher abundance of metabolites associated with caffeic and ferulic acid, which are part of a pathway biochemically distant from PD and HD biosynthesis (Yagi *et al*., 1977).

The metabolites umbelliferone, naringenin, and resveratrol displayed a positive correlation with PD and HD concentrations (Fig. 3G, D and Fig. S8A). This suggests that these metabolites might indeed be associated with PD and HD biosynthesis. Moreover, accessions characterized by high PD and/or HD levels exhibited elevated concentrations of these metabolites, along with Thn C (Fig. 3H). At the transcriptome level, genes of the enzymes CHS (p-coumaroyl CoA → naringenin chalcone), CHI (naringenin chalcone → naringenin), F3’5’H (dihydrokaempferol → dihydromyricetin), DFR (dihydromyricetin → leucodelphinidin), ROMT (resveratrol → pinostilbene → pterostilbene) and FOMT showed upregulation in high PD and/or HD accessions and downregulation in low PD/HD accessions (Fig. S8). By integrating the metabolite and transcriptome data, it can be inferred that accessions with high PD and/or HD levels exhibit enhanced flavonoid biosynthesis (involving metabolites derived from naringenin chalcone) and stilbene biosynthesis (involving metabolites derived from resveratrol) compared to low PD/HD accessions. Notably, previous studies have reported a diverse range of metabolites in Hydrangeas (Brown *et al*., 1964; Wellmann *et al*., 2022; Yoon *et al*., 2023; Yano *et al*., 2023). However, the present research successfully incorporates the underlying genetic changes, particularly in terms of metabolic regulation.

p-coumaroyltriacetic acid synthase (CTAS) plays a key role in the conversion of p-coumaroyl CoA to p-coumaroyltriacetic acid lactone (CTAL). Previous studies have hypothesized that CTAS, in conjunction with a cyclase and a ketoreductase, can facilitate the synthesis of hydrangeic acid. Subsequently, hydrangeic acid may undergo further modifications within the plant, ultimately leading to the formation of HD (Akiyama *et al*, 1999; Austin and Noel, 2003; Eckermann *et al*, 2003). In the current research, significantly higher transcript level of *CTAS* gene was observed in the accessions characterized by a high concentration of PD and/or HD, with the highest expression detected in the accessions containing both PD and HD (Fig. 7, Fig. S4-S6). Using transcriptome analysis and RT-qPCR, the present study revealed up-regulation of type III polyketide synthases (*PKS*), ketoreductase (*KR*), polyketide cyclase (*PKC*) and double bond reductase (*DBR*) in accessions with high PD and/or HD (Fig. 7, Fig. S4-S6).

Interestingly, *CTAS* gene was absent in *H. paniculata* and *PKC* together with *KR* showed elevated expression levels when all *H. macrophylla* accessions were compared with *H. paniculata*, where PD and HD are not detected (data not shown). This observation suggests that these genes probably play a central role in the biosynthesis of PD and HD. The abundance of these genes, particularly *KR* and *PKC*, could potentially form a multi-gene complex responsible for the synthesis of stilbene carboxylic acid (hydrangeic acid), in line with a hypothesis raised in previous research (Akiyama *et al*, 1999; Austin and Noel, 2003). A similar set of genes involving type *III PKS, KR* and *PKC* has been proposed to be responsible for the production of lunularic acid, another stilbene carboxylic acid, in *Cannabis sativa* L (Gülck and Møller, 2020). Several alternative pathways for the biosynthesis of PD and HD have been proposed in the literature, and an intriguing hypothesis revolves around the role of thunberginols in this biosynthetic pathway. Thunberginols and HD have been identified as catabolic products in the faeces of PD-fed rats (Yasuda *et al*, 2004 and Preusche *et al*, 2022). Furthermore, a proposed biosynthetic pathway starting from resveratrol to dihydroresveratrol, 5-hydroxy-lunularic acid, and Thn C has been proposed (Çiçek *et al*, 2018). In addition, a type III polyketide synthase (*PKS*) responsible for the conversion of dihydro-paracoumaroyl-CoA to dihydroresveratrol has been reported in *Cannabis sativa* L. (Boddington *et al*, 2022). In the current study, higher concentration of resveratrol was observed in accessions with high PD and/or HD levels compared to those with low PD/HD levels (Fig. 3G). The enzyme responsible for the downstream processing of resveratrol, ROMT, also showed increased expression levels in accessions with high PD and/or HD compared to low PD/HD accessions (Fig. 7A). At the same time, a higher relative abundance of Thn C was found in the accessions with high PD and/or HD levels compared to those with low PD/HD levels (Fig. 3H). However, it is noteworthy that resveratrol and Thn C were not detected in the *H. paniculata* accession (Fig. 3H). Most importantly, 5-hydroxy-lunularic acid, a metabolite involved in the same pathway, could not be evaluated due to the unavailability of a reference standard at the time of the study and the challenges associated with proper chromatographic separation. However, the presence of 5-hydroxylunularic acid and its closely related metabolites in hydrangeas has been mentioned in previous reports (Gorham, 1977 and Bojack *et al*, 2022). By incorporating the findings from the metabolomic and transcriptomic data in the present research, coupled with an extensive review of the relevant literature on the subject, a speculative pathway outlining the biosynthesis of PD has been devised and is shown in Fig. 8.

### Proposed model for phyllodulcin biosynthesis

Taken together, the results of the metabolomic and transcriptomic analyses indicate that in low PD/HD *H. macrophylla* accessions, there is a substantial diversion of metabolic flow away from dihydroisocoumarin biosynthesis. Conversely, in accessions containing PD, HD or both PD and HD, the pathway is directed towards dihydroisocoumarin biosynthesis. This clear differentiation between accessions based on their biochemical concentrations is evident in the PCA biplot (Fig. 4B) and metabolite heatmap (Fig. 4A, C). In particular, naringenin, phenylalanine, resveratrol, umbelliferone and, to some extent, esculin are responsible for the differences observed in the high PD accessions. Similarly, esculetin, scopolin, trans-cinnamic acid, scopoletin, caffeic acid and ferulic acid contribute to the characteristics of the low PD/HD accessions (Fig. 3). These pathway shifts, coupled with variations in the expression of key genes within the PPP, clearly show that the initial reactions for PD biosynthesis are the same as for many other secondary metabolites, but from p-coumaric acid, the branching point, three distinct pathways including HD, resveratrol and/or Thn C are most likely the main routes to PD biosynthesis, as demonstrated by the combination of our metabolomics and transcriptomics results in this study (Fig. 8). Which pathway is ultimately preferred needs to be explored by designing the pathway in a non-PD containing plant such as tobacco, transgenic approaches and/or feeding experiments with the direct labeled precursors of each pathway.

## Acknowledgements

We would like to thank Dr Jacob Ley and Esther-Corinna Schwarze (Symrise AG, Holzminden, Germany) for providing us the authentic external standards for PD and HD. We would also like to thank Thomas Becker (head of the hortensia company Kötterheinrich, Germany) for allowing us to grow and maintain the plants. We thank Nicole Schäfer, Melanie Ruff for their excellent assistance in harvesting the plant material and preparing the samples. We acknowledge the help and support of our horticultural team led by Enk Geyer and his colleagues in growing and maintaining the plants at IPK.

## Authorship contribution

G.P.S. and M.R.H. conceived the study and designed the experiments. G.P.S and M.R.H. drafted and critically revised the manuscript. F.E. grew, harvested and selected the plants for all experiments. A.H. contributed to the analysis and interpretation of the transcriptomic data. N.v.W. contributed with scientific suggestions, revision of the manuscript and data interpretation. We also thank Dr. Drescher and Sarah Maitri Bastian from AiF for their excellent support during the project.

## Competing interests

The authors declare that the research was conducted in the absence of any financial interests/personal relationships that could be considered as a potential conflict of interest.

## Funding

This study was funded by Federal Ministry for Economic Affairs and Energy (BMWi) coordinated by the German Federation of Industrial Research Associations (AiF).

## Data availability

The RNA sequencing dataset is available on ENA Browser-European Nucleotide Archive.

## Notes

### Competing Interest Statement

The authors have declared no competing interest.

