## Supplementary Figures for "Elucidation and functional characterization of natural sweetener biosynthesis in hortensia species"

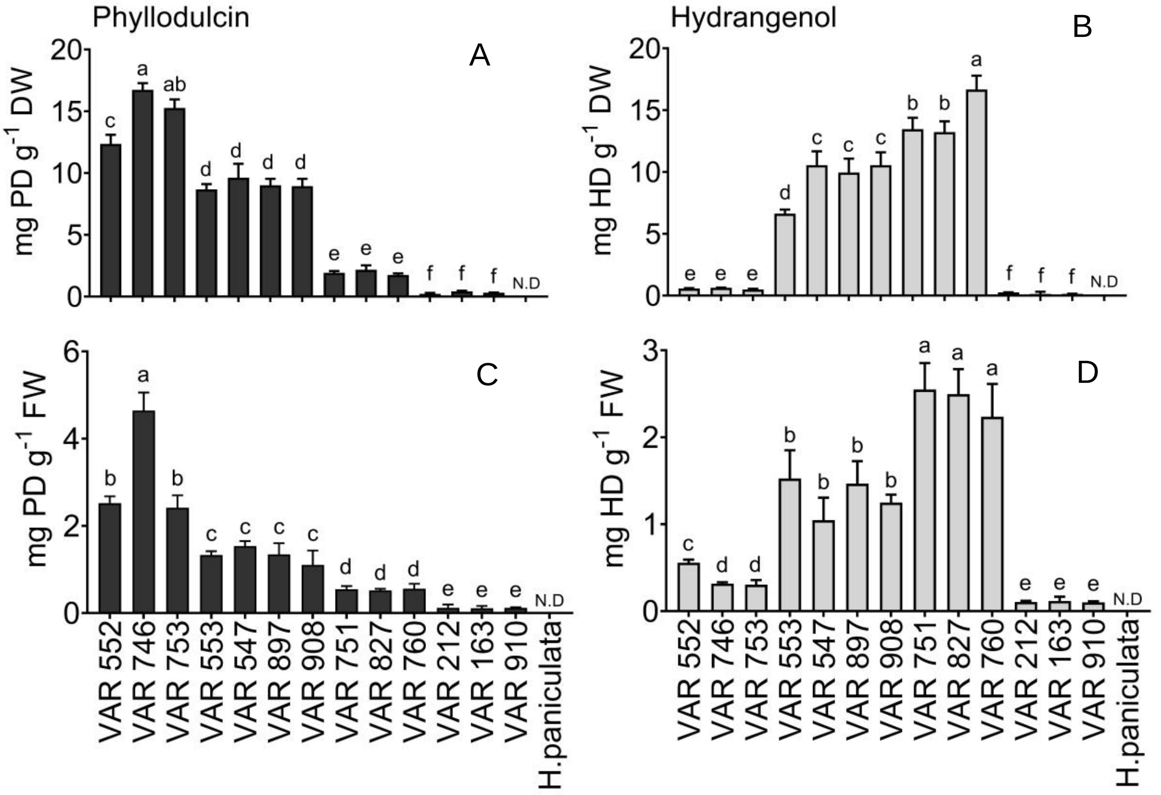


**Fig. S1.** Comparison between fresh and dried leaf tissues for phyllodulcin (PD) and hydrangenol (HD) concentrations in selected Hydrangea accessions. (A) PD and (B) HD concentrations in dried leaves. (C) PD and (C) HD concentrations in freshly harvested leaves. Analysis was performed on fully expanded young upper leaves. Bars represent mean + SE (n=6). Not detected (N.D.) indicates the absence of the corresponding metabolite. Different letters indicate significant differences between accessions according to one-way ANOVA followed by post-hoc Tukey's test ($p<0.05$).

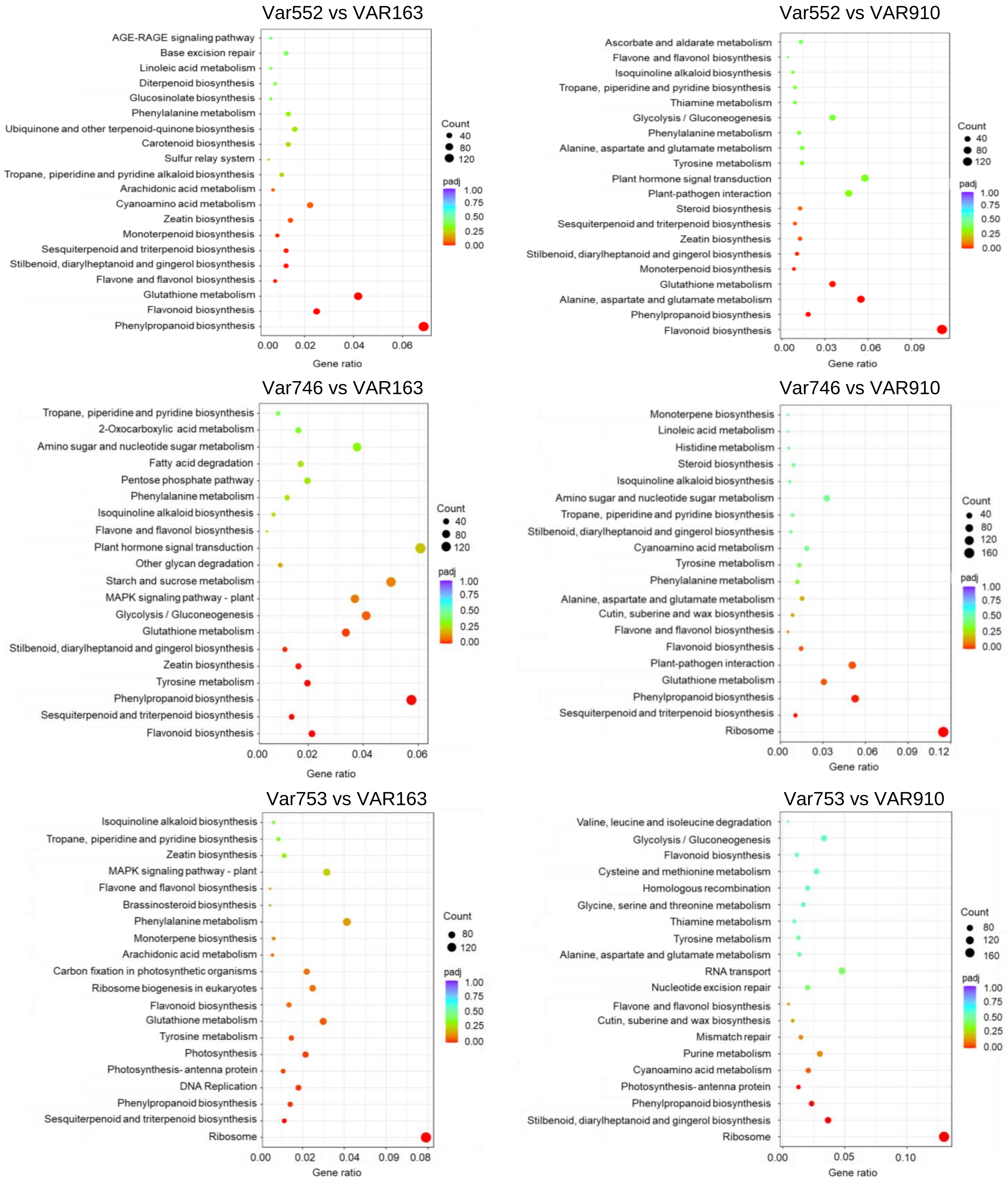


**Fig. S2.** The assignment of genes to different pathways in high PD (VAR552, VAR746, VAR753) and low PD (VAR163, VAR910) containing accessions of H. macrophylla. (A-F) Dot plot of the KEGG enrichment analysis showing the gene ratio (the percentage of total DEGs) assigned to the top 20 pathways in the study group. The dot size represents the number of genes and the colour of the dot is based on the p-value adjusted to the sample distribution (padj value) and indicates the significance of pathway enrichment. KEGG annotates genes at the pathway level.

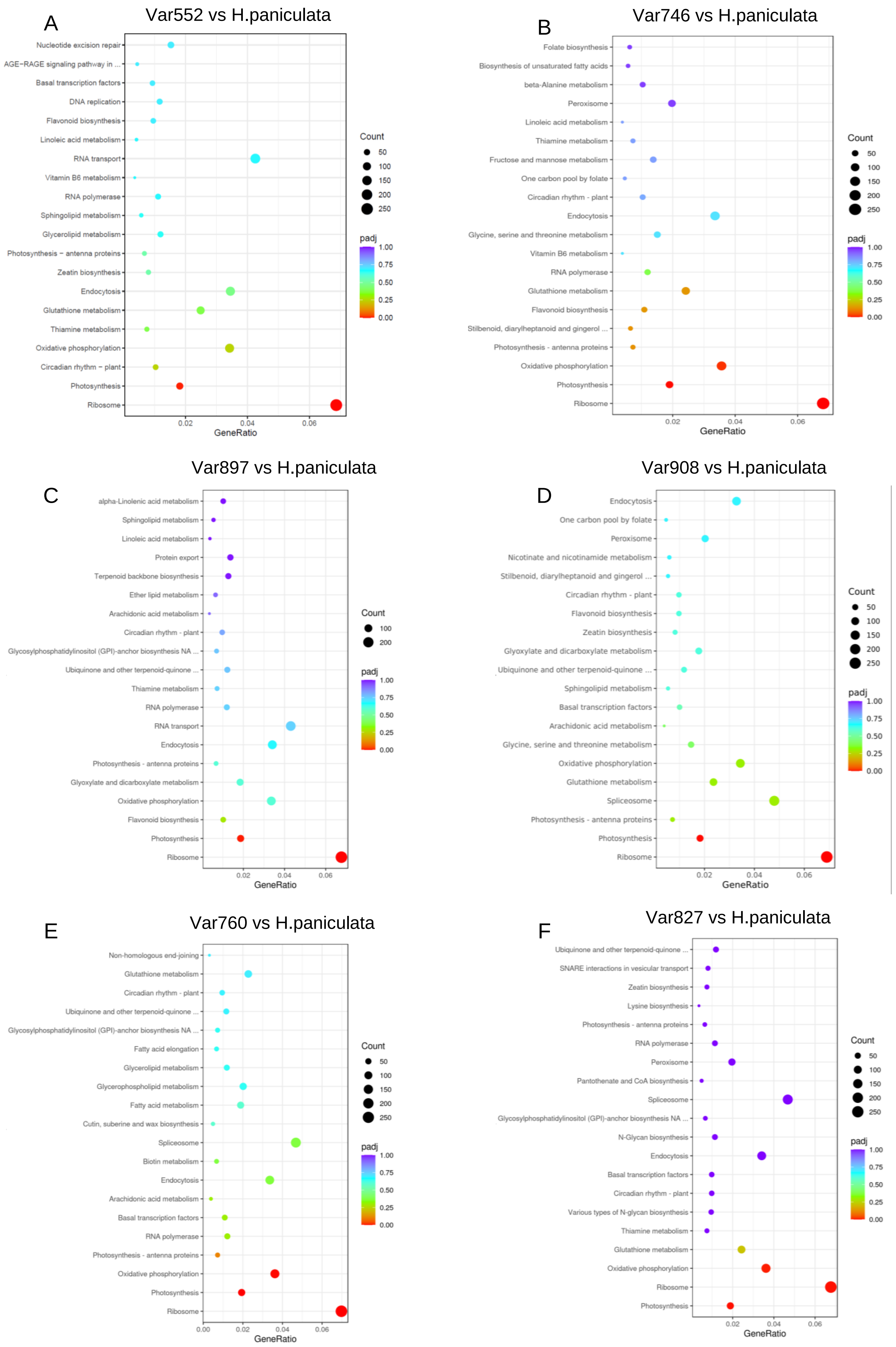
A
Var552 vs H.paniculata
Nucleotide excision repair
AGE−RAGE signaling pathway in ...
Basal transcription factors
DNA replication
Flavonoid biosynthesis
Linoleic acid metabolism
RNA transport
Vitamin B6 metabolism
RNA polymerase
Sphingolipid metabolism
Glycerolipid metabolism
Photosynthesis − antenna proteins
Zeatin biosynthesis
Endocytosis
Glutathione metabolism
Thiamine metabolism
Oxidative phosphorylation
Circadian rhythm − plant
Photosynthesis
Ribosome
0.02
0.04
0.06
GeneRatio
Count
50
100
150
200
250
padj
1.00
0.75
0.50
0.25
0.00
B
Var746 vs H.paniculata
Folate biosynthesis
Biosynthesis of unsaturated fatty acids
beta-Alanine metabolism
Peroxisome
Linoleic acid metabolism
Thiamine metabolism
Fructose and mannose metabolism
One carbon pool by folate
Circadian rhythm - plant
Endocytosis
Glycine, serine and threonine metabolism
Vitamin B6 metabolism
RNA polymerase
Glutathione metabolism
Flavonoid biosynthesis
Stilbenoid, diarylheptanoid and gingerol ...
Photosynthesis - antenna proteins
Oxidative phosphorylation
Photosynthesis
Ribosome
0.02
0.04
0.06
GeneRatio
Count
50
100
150
200
250
padj
1.00
0.75
0.50
0.25
0.00
C
Var897 vs H.paniculata
alpha-Linolenic acid metabolism
Sphingolipid metabolism
Linoleic acid metabolism
Protein export
Terpenoid backbone biosynthesis
Ether lipid metabolism
Arachidonic acid metabolism
Circadian rhythm - plant
Glycosylphosphatidylinositol (GPI)-anchor biosynthesis NA ...
Ubiquinone and other terpenoid-quinone ...
Thiamine metabolism
RNA polymerase
RNA transport
Endocytosis
Photosynthesis - antenna proteins
Glyoxylate and dicarboxylate metabolism
Oxidative phosphorylation
Flavonoid biosynthesis
Photosynthesis
Ribosome
0.02
0.04
0.06
GeneRatio
Count
100
200
padj
1.00
0.75
0.50
0.25
0.00
D
Var908 vs H.paniculata
Endocytosis
One carbon pool by folate
Peroxisome
Nicotinate and nicotinamide metabolism
Stilbenoid, diarylheptanoid and gingerol ...
Circadian rhythm - plant
Flavonoid biosynthesis
Zeatin biosynthesis
Glyoxylate and dicarboxylate metabolism
Ubiquinone and other terpenoid-quinone ...
Sphingolipid metabolism
Basal transcription factors
Arachidonic acid metabolism
Glycine, serine and threonine metabolism
Oxidative phosphorylation
Glutathione metabolism
Spliceosome
Photosynthesis - antenna proteins
Photosynthesis
Ribosome
0.02
0.04
0.06
GeneRatio
Count
50
100
150
200
250
padj
1.00
0.75
0.50
0.25
0.00
E
Var760 vs H.paniculata
Non-homologous end-joining
Glutathione metabolism
Circadian rhythm - plant
Ubiquinone and other terpenoid-quinone ...
Glycosylphosphatidylinositol (GPI)-anchor biosynthesis NA ...
Fatty acid elongation
Glycerolipid metabolism
Glycerophospholipid metabolism
Fatty acid metabolism
Cutin, suberine and wax biosynthesis
Spliceosome
Biotin metabolism
Endocytosis
Arachidonic acid metabolism
Basal transcription factors
RNA polymerase
Photosynthesis - antenna proteins
Oxidative phosphorylation
Photosynthesis
Ribosome
0.00
0.02
0.04
0.06
GeneRatio
Count
50
100
150
200
250
padj
1.00
0.75
0.50
0.25
0.00
F
Var827 vs H.paniculata
Ubiquinone and other terpenoid-quinone ...
SNARE interactions in vesicular transport
Zeatin biosynthesis
Lysine biosynthesis
Photosynthesis - antenna proteins
RNA polymerase
Peroxisome
Pantothenate and CoA biosynthesis
Spliceosome
Glycosylphosphatidylinositol (GPI)-anchor biosynthesis NA ...
N-Glycan biosynthesis
Endocytosis
Basal transcription factors
Circadian rhythm - plant
Various types of N-glycan biosynthesis
Thiamine metabolism
Glutathione metabolism
Oxidative phosphorylation
Ribosome
Photosynthesis
0.02
0.04
0.06
GeneRatio
Count
50
100
150
200
250
padj
1.00
0.75
0.50
0.25
0.00

**Fig. S3.** The assignment of genes to different pathways in high PD (A, B, VAR552, VAR746), high PD/HD (C, D, VAR897, VAR908) or high HD (E, F, VAR760, VAR827) and the negative control H.paniculate accessions of H. macrophylla. (A-F) Dot plot of the KEGG enrichment analysis showing the gene ratio (the percentage of total DEGs) assigned to the top 20 pathways in the study group. The dot size represents the number of genes and the colour of the dot is based on the p-value adjusted to the sample distribution (padj value) and indicates the significance of pathway enrichment. KEGG annotates genes at the pathway level.

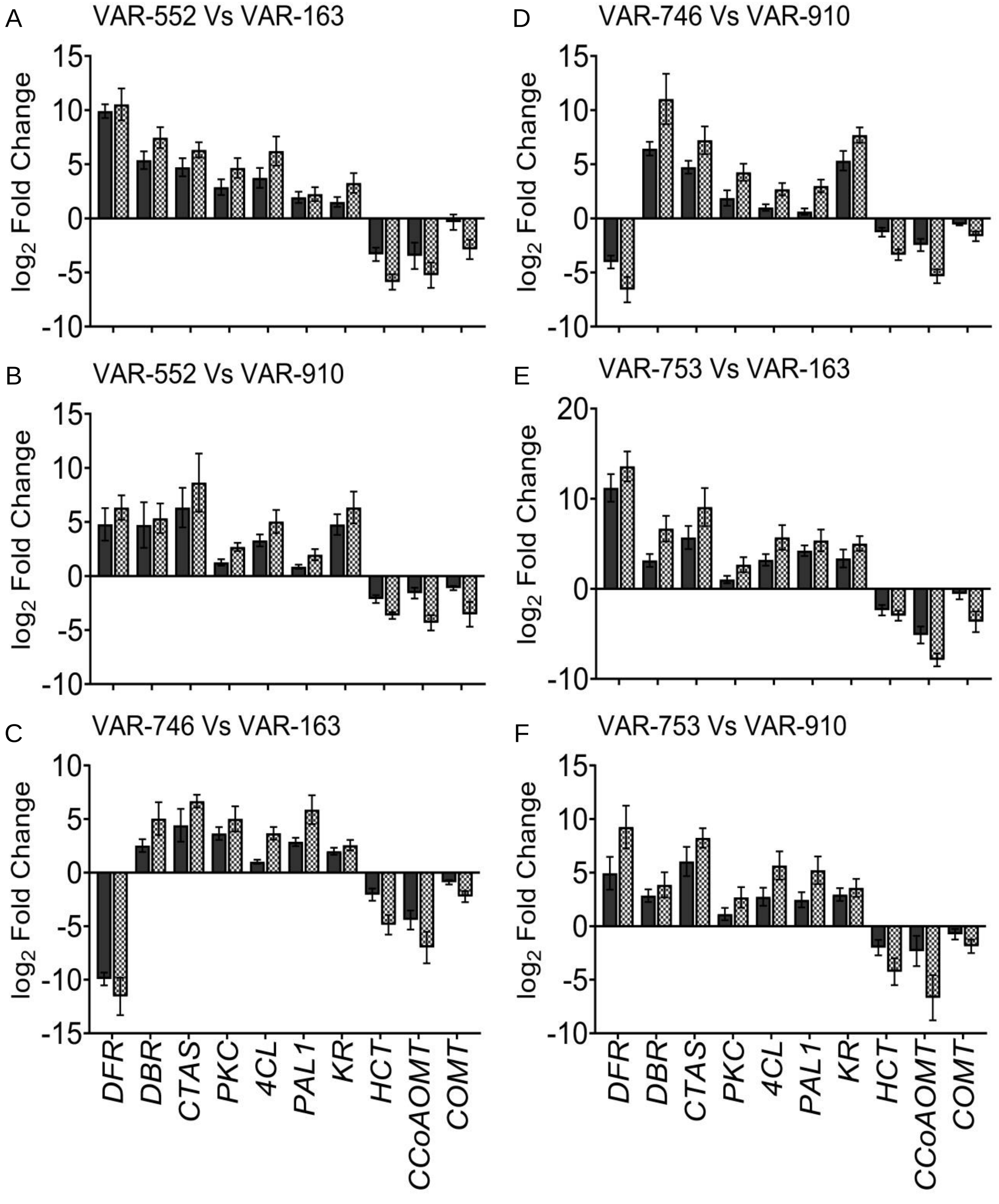


**Fig. S4.** Validation of the expression of candidate genes by means of quantitative real-time PCR (qRT-PCR). (A-F) The expression of genes involved in the metabolic pathways associated with PD biosynthesis while comparing accessions (A) VAR-552 and VAR-163, (B) VAR-552 and VAR-910, (C) VAR-746 and VAR-163 and (D) VAR-746 and VAR-910, (E) VAR-753 and VAR-163 and (F) VAR-753 and VAR-910. Ten representative genes namely DFR, DBR, CTAS, PKC, 4CL, PAL1, KR, HCT, CCoAOMT and COMT were selected to validate the expression of genes obtained from RNAseq using RT-qPCR. Bars indicate mean ± SE (n=3) of log2 fold-change obtained from the respective experiments. Positive values represent upregulation and negative values represent downregulation. The gene GADPH was used as an internal reference gene. Black bars show log2 fold change of RNASeq data and hatched bars show the log2 fold change of qRT-PCR.

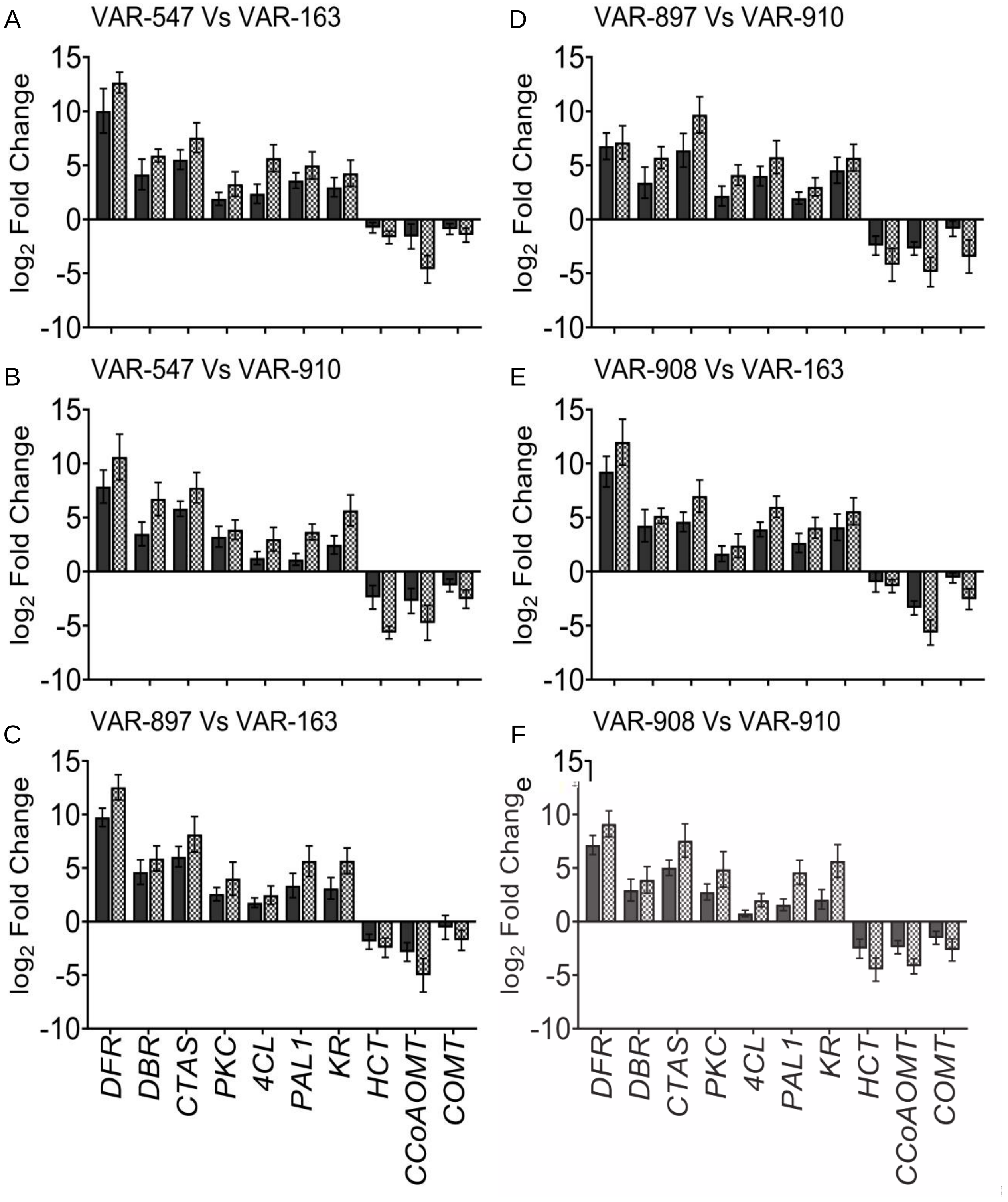


**Fig. S5.** Validation of the expression of candidate genes by means of quantitative real-time PCR (qRT-PCR). (A-F) The expression of genes involved in the metabolic pathways associated with PD biosynthesis while comparing accessions (A) VAR-547 and VAR-163, (B) VAR-547 and VAR-910, (C) VAR-897 and VAR-163 and (D) VAR-897 and VAR-910, (E) VAR-908 and VAR-163 and (F) VAR-908 and VAR-910. Ten representative genes namely DFR, DBR, CTAS, PKC, 4CL, PAL1, KR, HCT, CCoAOMT and COMT were selected to validate the expression of genes obtained from RNAseq using RT-qPCR. Bars indicate mean ± SE (n=3) of log2 fold-change obtained from the respective experiments. Positive values represent upregulation and negative values represent downregulation. The gene GADPH was used as an internal reference gene. Black bars show log2 fold change of RNASeq data and hatched bars show the log2 fold change of qRT-PCR.

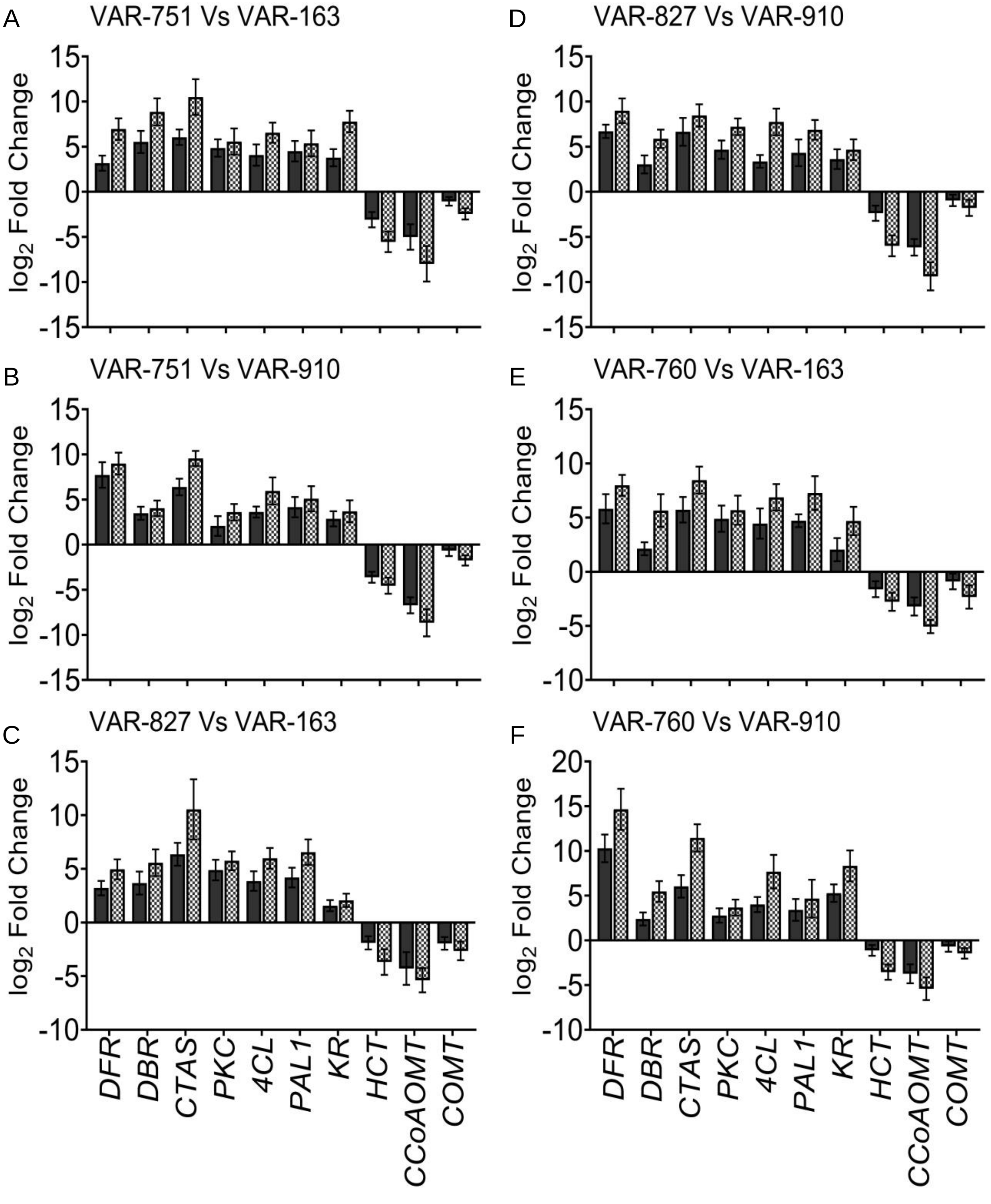


**Fig. S6.** Validation of the expression of candidate genes by means of quantitative real-time PCR (qRT-PCR). (A-F) The expression of genes involved in the metabolic pathways associated with PD biosynthesis while comparing accessions (A) VAR-751 and VAR-163, (B) VAR-751 and VAR-910, (C) VAR-827 and VAR-163 and (D) VAR-827 and VAR-910, (E) VAR-760 and VAR-163 and (F) VAR-760 and VAR-910. Ten representative genes namely DFR, DBR, CTAS, PKC, 4CL, PAL1, KR, HCT, CCoAOMT and COMT were selected to validate the expression of genes obtained from RNAseq using RT-qPCR. Bars indicate mean ± SE (n=3) of log2 fold-change obtained from the respective experiments. Positive values represent upregulation and negative values represent downregulation. The gene GADPH was used as an internal reference gene. Black bars show log2 fold change of RNASeq data and hatched bars show the log2 fold change of qRT-PCR.

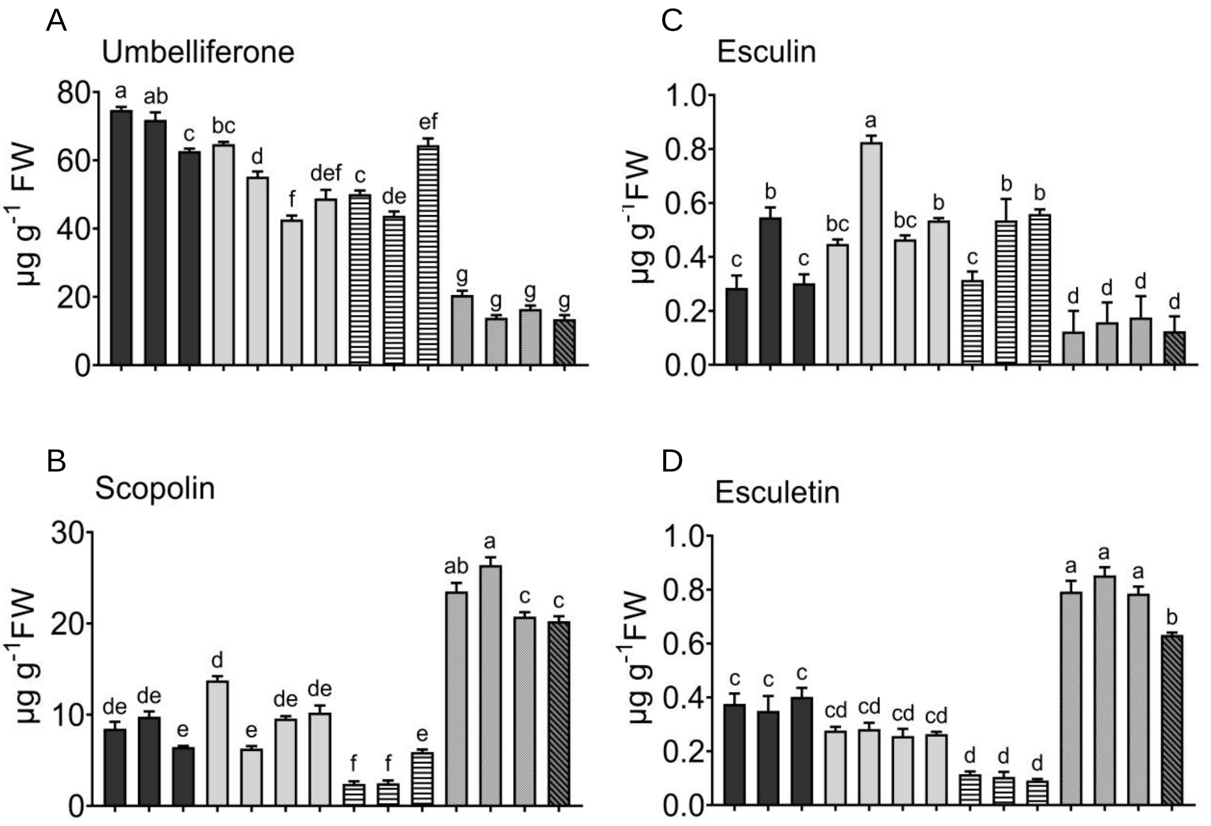


**Fig. S7. Levels of coumarins across various Hydrangea accessions**. (A) umbelliferone, (B) scopolin, (C) esculin, and (D) esculetin measured in 14 selected accessions of Hydrangea. Analysis was performed on freshly harvested, fully expanded leaves in plants grown for 75 days. Bars represent the mean of 6 independent biological replicates (n=6) and standard error. Not detected (N.D.) indicates the absence of the corresponding metabolite. Dark bars show accessions with high PD, light grey bars accessions with high PD/HD, hatched bars accessions with high HD, dark grey bars accessions with low PD/HD and dark hatched bars H. paniculate as negative control. Different letters denote significant differences among varieties according to one-way ANOVA and post-hoc Tukey’s test ($p<0.05$).

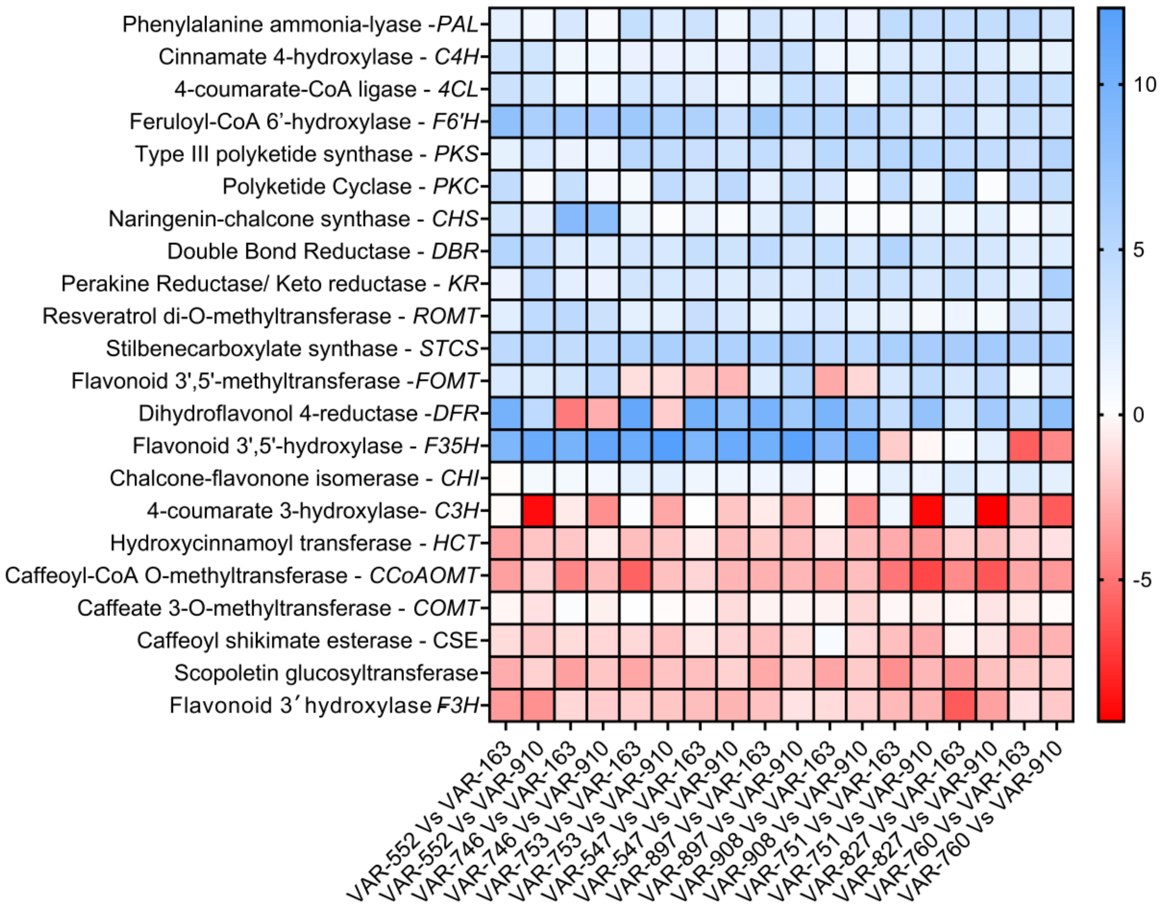


**Fig. S8.** Expression patterns of module-specific genes involved in phenylpropanoid pathway and related pathways in H. macrophylla accessions. DEGs were selected based on module-trait relationships derived from WGCNA, with each gene in a module correlating with individual metabolite concentrations (padj<0.05). Each black square represents the normalized log-fold2 change values of the specific gene in each study group. The comparison was made between a high PD and/or HD accession and low PD/HD accessions (VAR-163 and VAR-910). Dark blue represents the highest relative expression level of the gene and dark red represents the lowest relative expression level of the gene.

**Table S1**. correlation analysis between identified genes and metabolites using WGCNA. PD: Dihydroisocoumarin, HD: hydrangenol. For gene names refer to the manuscript.

| gene no | Module colour | Gene count | Gene name Up-regulated | Gene name down-regulated | Positive correlation with metabolites |
|---|---|---|---|---|---|
| 1 | darkolivegreen | 5065 | PAL1, 4CL1, F6'H2, PKS, PKC | F3'H | PD, p-coumaric acid, trans-cinnamic acid |
| 2 | coral2 | 4153 | | | |
| 3 | darkmagenta | 3294 | | | |
| 4 | darkviolet | 2508 | | COMT | |
| 5 | green | 1907 | | | HD |
| 6 | bisque4 | 1656 | CTAS, ROMT, CHS, CHI | | PD, p-coumaric acid, resveratrol |
| 7 | indianred4 | 923 | | | scopoletin |
| 8 | floralwhite | 906 | 4CL1, F6'H2, DFR | | PD, HD, phenylalanine, p-coumaric acid, umbelliferone, naringenin |
| 9 | magenta | 768 | | | HD |
| 10 | brown2 | 553 | | HCT, C3H, CCoAOMT, SGT | Caffeic acid, ferrulic acid, scopoletin |
| 11 | darkturquoise | 432 | | | HD, resveratrol |
| 12 | navajowhite1 | 379 | F6'H2 | CSE | trans-cinnamic acid, caffeic acid, ferulic acid, esculetin, scopoletin |
| 13 | lightsteelblue | 172 | C4H | | PD, phenylalanine, p-coumaric acid |
| 14 | antiquewhite | 123 | | | P-coumaric acid |
| 15 | thistle | 73 | | | |
| 16 | grey | 68 | | | |
| | **TOTAL** | **22,980** | | | |

**Table S2**. List of qRT-PCR primers used for validation of identified differentially expressed genes via RNA sequencing

| Gene | Forward primer 5'→3' | Reverse primer 5'→3' |
|---|---|---|
| Glyceraldehyde-3-phosphate dehydrogenase - *GADPH* | GGCTGAGACTGGAGCGGAAT | CAAACATGGGGGCGTCTTTGC |
| Double Bond Reductase - *DBR* | ACCGTTGTGCCTTACATCAG | ACGGCCACTAAAGAGTCCAA |
| Dihydroflavonol 4-reductase - *DFR* | ATGTTGGACGCCTCTCCATC | CACAAGTCCACACCCAAGAG |
| p-Coumaroyl tri acetic acid synthase - *CTAS* | CAGGCAAAAGTGGGTCTGAA | GAAAAACACACATGCGCTGG |
| Polyketide Cyclase - PKC | TGCAGCCCATTCCAAATCAG | TGATGAAGGACCTTTCGGGA |
| Phenylalanine ammonia-lyase 1 – *PAL1* | ATCAGAGAGTGCCGGTCTTT | ACCGGACTTTCTCTCCTGTC |
| 4-coumarate CoA ligase – *4CL* | TGGTTCCAAGATCTCCGAGG | GAGCTTTTGGGATTGCGTCT |
| Keto reductase - *KR* | GGTAGCCCTGAATATGTGCG | GTCGACACGATGTGGGTAGT |
| Hydroxycinnamoyl transferase - *HCT* | AGATGGACTTTCAGCCCTCC | GAGGATGGTCCGATCGATGA |
| Caffeoyl-CoA 3-O-methyltransferase - *CCoAOMT* | GGCATGGAGCACAAGATCAA | GGGCTTGTCAGCGTCAATAA |
| Caffeic acid O-methyltransferase - *COMT* | GTGTCGATCGGAAGGCCATA | ATGCGACAGCACAAGGTAAC |

**Table S3a. List of *H. macrophylla* accessions used for primary screening.**
Green filled cells represent the 13 accessions selected for experiments.

| | | |
|---|---|---|
| VAR-0908_000 (VAR 908) | VAR-0553_001 (VAR 553) | VAR-0897_000 (VAR 897) |
| VAR-0553_002 | VAR-0552_000 (VAR 552) | VAR-0552_001 |
| VAR-0828_000 | VAR-0766_001 | VAR-0158_000 |
| VAR-0547_000 (VAR-547) | VAR-0542_000 | VAR-0536_000 |
| VAR-0749_000 | VAR-0556_000 | VAR-0912_000 |
| VAR-0765_000 | VAR-0906_000 | VAR-0583_000 |
| VAR-0556_001 | VAR-0836_000 | VAR-0571_000 |
| VAR-0844_000 | VAR-0535_000 | VAR-0576_000 |
| VAR-0879_001 | VAR-0768_000 | VAR-0827_001 |
| VAR-0827_000 (VAR 827) | VAR-0556_002 | VAR-0539_000 |
| VAR-0582_000 | VAR-0562_000 | VAR-0561_000 |
| VAR-0751_000 (VAR 751) | VAR-0580_000 | VAR-0560_000 |
| VAR-0574_000 | VAR-0260_002 | VAR-0763_000 |
| VAR-0550_000 | VAR-0575_000 | VAR-0561_001 |
| VAR-0750_000 | VAR-0747_000 | VAR-0573_000 |
| VAR-0566_000 | VAR-0921_000 | VAR-0446_000 |
| VAR-0846_000 | VAR-0547_002 | VAR-0567_000 |
| VAR-0555_000 | VAR-0564_000 | VAR-0706_000 |
| VAR-0915_000 | VAR-0132_002 | VAR-0438_000 |
| VAR-0746_000 (VAR 746) | VAR-0768_001 | VAR-0569_000 |
| VAR-0750_001 | VAR-0212_001 | VAR-0843_000 |
| VAR-0009_000 | VAR-0922_001 | VAR-0856_000 |
| VAR-0097_000 | VAR-0837_000 | VAR-0452_000 |
| VAR-0146_000 | VAR-0440_000 | VAR-0163_000 (VAR 163) |
| VAR-0579_000 | VAR-0543_000 | VAR-0823_000 |
| VAR-0771_000 | VAR-0260_000 | VAR-0538_000 |
| VAR-0758_000 | VAR-0554_000 | VAR-0913_000 |
| VAR-0551_000 | VAR-0754_000 | VAR-0290_000 |
| VAR-0755_000 | VAR-0745_000 | VAR-0565_000 |

**Table S3b. List of *H. macrophylla* accessions used for primary screening.** Green filled cells represent the 13 accessions selected for experiments.

| | | |
|---|---|---|
| VAR-0436_000 | VAR-0919_000 | VAR-0762_000 |
| VAR-0347_000 | VAR-0893_000 | VAR-0118_000 |
| VAR-0756_000 | VAR-0707_000 | VAR-0904_000 |
| VAR-0922_002 | VAR-0578_000 | VAR-0255_000 |
| VAR-0548_000 | VAR-0581_000 | VAR-0808_000 |
| VAR-0616_000 | VAR-0708_000 | VAR-0808_001 |
| VAR-0491_000 | VAR-0905_000 | VAR-0069_000 |
| VAR-0348_000 | VAR-0258_000 | VAR-0753_001 (VAR 753) |
| VAR-0142_000 | VAR-0766_000 | VAR-0856_001 |
| VAR-0540_000 | VAR-0890_000 | VAR-0145_000 |
| VAR-0256_000 | VAR-0010_000 | VAR-0704_000 |
| VAR-0741_000 | VAR-0544_000 | VAR-0549_000 |
| VAR-0559_000 | VAR-0068_000 | VAR-0577_000 |
| VAR-0572_000 | VAR-0584_000 | VAR-0364_000 |
| VAR-0477_000 | VAR-0132_000 | VAR-0769_000 |
| VAR-0879_002 | VAR-0757_000 | VAR-0568_000 |
| VAR-0439_000 | VAR-0537_000 | VAR-0615_000 |
| VAR-0823_001 | VAR-0259_000 | VAR-0811_000 |
| VAR-0492_000 | VAR-0918_000 | VAR-0748_000 |
| VAR-0761_000 | VAR-0907_000 | VAR-0770_000 |
| VAR-0131_000 | VAR-0770_001 | VAR-0570_000 |
| VAR-0493_000 | VAR-0758_002 | VAR-0447_000 |
| VAR-0039_000 | VAR-0418_000 | VAR-0920_000 |
| VAR-0856_002 | VAR-0772_000 | VAR-0451_000 |
| VAR-0744_000 | VAR-0835_000 | VAR-0760_000 (VAR 760) |
| VAR-0212_000 (VAR 212) | VAR-0917_000 | VAR-0879_003 |
| VAR-0163_001 | VAR-0437_000 | VAR-0901_000 |
| VAR-0891_000 | VAR-0910_000 (VAR 910) | VAR-0894_000 |
| VAR-0898_000 | VAR-0896_000 | VAR-0857_000 |
| VAR-0758_001 | VAR-0900_000 | VAR-0119_000 |
| VAR-0824_001 | VAR-0911_000 | VAR-0899_000 |
| VAR-0895_000 | VAR-0563_000 | |

**Supplementary protocol S1. RNA isolation, sequencing and analysis**

To isolate total RNA from the plant tissues, the Spectrum™ Plant Total RNA Kit, obtained from Sigma Aldrich, was employed following the manufacturer's guidelines, and it was found to be highly effective for isolating RNA from Hydrangea tissues. Total RNA was isolated and analyzed from three replicates of 14 selected accessions, each with varying levels of PD and HD concentration. The quality of RNA was assessed using NanoDrop™ 2000c spectrophotometry from Thermo Scientific™, with a 10 µL volume collected and used for sequencing. RNA integrity was further evaluated using the RNA Nano 6000 Assay Kit on the Bioanalyzer 2100 system by Agilent Technologies in California, USA. For cDNA library preparation and transcriptome sequencing, Novogene conducted the procedures using the NEBNext® UltraTM RNA Library Prep Kit for Illumina® (NEB, USA). Initially, mRNA was purified from total RNA using poly-T oligo-attached magnetic beads, followed by fragmentation. Subsequently, the first and second strands of cDNA were synthesized using a random hexamer primer and M-MuLV Reverse Transcriptase (RNase H-) along with DNA Polymerase I and RNase H, respectively. cDNA fragments of lengths between 370 to 420 bp were selected after purifying library fragments using the AMPure XP system from Beckman Coulter in Beverly, USA. PCR was then carried out with Phusion High-Fidelity DNA polymerase, Universal PCR primers, and Index (X) Primer. The quality of the libraries was assessed using the Agilent Bioanalyzer 2100 system. Upon library preparation, 150 bp pair end reads were generated after clustering. The reference genome and gene model annotation files were downloaded directly from https://plantgarden.jp. Indexing of the reference genome and read alignment to the reference genome were performed using Hisat2 v2.0.5. The mapped reads from each sample were assembled using StringTie (v1.3.3b) (Pertea et al, 2015) in a reference-based approach. FeatureCounts v1.5.0-p3 was utilized to count the number of reads mapped to each gene. Differential expression analysis was conducted using the DESeq2 R package (v1.20.0). The resulting p-values were adjusted using the Benjamini and Hochberg’s approach to control the False Discovery Rate (FDR), represented as padj value. Significantly differentially expressed genes were identified with a corrected p-value of 0.05 and an absolute fold change of 2 as the threshold. To gain insights into the functional significance of differentially expressed genes, Gene Ontology (GO) enrichment analysis was performed using the clusterProfiler R package, with GO terms exhibiting corrected p-values below 0.05 considered significantly enriched by the differentially expressed genes. Additionally, KEGG (Kyoto Encyclopedia of Genes and Genomes) pathway enrichment analysis was conducted to assign each gene to its respective pathways.

**Supplementary protocol S2. Detailed description of qRT-PCR method**

The RevertAid First Strand cDNA Synthesis Kit from Thermo Scientific was utilized to perform cDNA synthesis, and 0.3 µg of total RNA was employed as the starting material. The synthesis was initiated using oligo(dT) primers. The primers for the quantitative real-time polymerase chain reaction (qRT-PCR) were created through primer design software Primer3, and they were subsequently synthesized by the company Metabion (Germany). The list of primers designed for validation with qRT-PCR are represented in Table S1. Specific criteria during the primer design process included achieving a melting temperature (Tm) within the range of 60±1°C, designing primers with a length between 18 and 25 base pairs, preferably located close to the 3'-end of the target sequence, and ensuring a GC content between 40% and 60%. This strategy aimed to generate PCR products that are unique and relatively short, ranging from 60 to 150 base pairs in length. Gene expression level was employed using qPCR (TouchTM system, Bio-Rad. For the qPCR reactions, the iQ SYBR Green Supermix from Bio-Rad (Hercules, CA, USA) was used. The samples underwent examination in triplicates, following this procedure: an initial activation cycle was carried out for 3 minutes at 95°C, accompanied by 40 amplification cycles, with each cycle comprising 15 seconds at 95°C and then 30 seconds at either 58°C or 60°C. Subsequently, a single melting curve cycle was executed, spanning from 65°C to 95°C, with increments of 0.5°C every 5 seconds. The Ct (Cycle threshold) values obtained were exported from the CFX Software (Bio-Rad Laboratories) and were employed to calculate the efficiency of PCR amplification. Normalization factors were calculated using glyceraldehyde-3-phosphate dehydrogenase (GAPDH) as reference gene. PCR amplification efficiency was determined following the guidelines outlined in prior instructions (Bustin et al, 2009). Only experiments exhibiting an efficiency falling within the range of 90% to 110% were considered. Normalization factors were calculated using geNORM (Vandesompele et al, 2002). The expression levels of genes were represented as fold change.
